# Evolution and connectivity influence the persistence and recovery of coral reefs under climate change in the Caribbean, Southwest Pacific, and Coral Triangle

**DOI:** 10.1101/2021.02.04.429453

**Authors:** Lisa C. McManus, Daniel L. Forrest, Edward W. Tekwa, Daniel E. Schindler, Madhavi A. Colton, Michael M. Webster, Timothy E. Essington, Stephen R. Palumbi, Peter J. Mumby, Malin L. Pinsky

## Abstract

Corals are experiencing unprecedented decline from climate change-induced mass bleaching events. Dispersal not only contributes to coral reef persistence through demographic rescue but can also hinder or facilitate evolutionary adaptation. Locations of reefs that are likely to survive future warming therefore remain largely unknown, particularly within the context of both ecological and evolutionary processes across complex seascapes that differ in temperature range, strength of connectivity, network size, and other characteristics. Here, we used eco-evolutionary simulations to examine coral adaptation to warming across reef networks in the Caribbean, the Southwest Pacific, and the Coral Triangle. We assessed the factors associated with coral persistence in multiple reef systems to understand which results are general and which are sensitive to particular geographic contexts. We found that evolution can be critical in preventing extinction and facilitating the long-term recovery of coral communities in all regions. Furthermore, the strength of immigration to a reef (destination strength) and current sea surface temperature robustly predicted reef persistence across all reef networks and across temperature projections. However, we found higher initial coral cover, slower recovery, and more evolutionary lag in the Coral Triangle, which has a greater number of reefs and more larval settlement than the other regions. We also found the lowest projected future coral cover in the Caribbean. These findings suggest that coral reef persistence depends on ecology, evolution, and habitat network characteristics, and that, under an emissions stabilization scenario (RCP 4.5), recovery may be possible over multiple centuries.

## Introduction

Rapidly increasing temperatures are threatening coral populations around the world through mass bleaching events that have increased in frequency and severity in recent decades (Terry P. Hughes et al., 2018). Though projections of coral persistence into the future are often dire (Hoegh-Guldberg et al., 2017), recent work highlights potential evolutionary mechanisms that may facilitate coral adaptation to warming waters (Bay et al., 2017; Kleypas et al., 2016). This is particularly relevant in the context of networks or metapopulations because each reef’s adaptive capacity is constrained by the balance between selection and migration (Lenormand, 2002). Subpopulations experience selection to adapt to their local environment, yet can also receive immigrants adapted to different environments. While there is increasing recognition for the contribution of evolution to coral persistence under future conditions (Logan et al., 2014; Matz et al., 2018, 2020a; Walsworth et al., 2019), there is less understanding of the interactions between evolutionary potential and reef characteristics on coral survival within regional-scale reef networks.

From a demographic perspective, larval dispersal links coral reefs within a network. Connectivity matrices describe the probabilities of viable larvae reaching one reef from another through dispersal and are typically generated from ocean circulation models (Kool et al., 2011; Treml et al., 2008; Watson et al., 2010) or population genetics data (Davies et al., 2015; Galindo et al., 2010; Matz et al., 2018). An important metric of connectivity is destination strength, which is the sum of incoming connection probabilities into a particular patch and is associated with a greater probability of reef survival (McManus et al., 2020). While modeling studies have explored the ecological importance of particular sites within a metapopulation (Kininmonth et al., 2019; Watson et al., 2011), few papers examine the combined ecological and evolutionary consequences of connectivity that are likely to be important across metapopulations (but see Matz et al., 2020).

In addition, the relative importance of connectivity to coral persistence as compared to local factors like baseline temperature and warming remains unclear. For example, bleaching records suggest that reefs that have experienced warm temperatures in the past are less likely to bleach in the future (Guest et al., 2012). Because temperatures are rising, coral populations in cooler conditions will likely benefit from receiving larvae that are pre-adapted to warmer environments, while relatively warm reefs are susceptible to the arrival of maladapted larvae (Norberg et al., 2012). In addition, temperature changes are likely to vary from one location to another (e.g., Ban et al., 2012). Generally, populations that experience a faster rate of environmental change are less likely to adapt (Lindsey et al., 2013), suggesting variation in extinction susceptibilities across reef networks. Due to evidence of thermal adaptation and heritability of heat tolerance in corals (Dixon et al., 2015; Dziedzic et al., 2019a; Kirk et al., 2018a), metrics that quantify the relative temperature of a patch and overall temperature change may be consequential determinants of individual reef persistence. In fact, in a recent modeling study of corals in the Indo-West Pacific, both the proportion of recruits immigrating from warmer locations and the present-day temperature were found to be useful factors in determining corals’ projected adaptive response (Matz et al., 2020a). Therefore, from an evolutionary perspective, larval dispersal facilitates the exchange of traits across a network.

Climate change is impacting reefs across the world, but the rates of temperature change (McClanahan et al., 2019) and the structure of dispersal networks (Wood et al., 2014) differ substantially across regions. While most modeling work addressing coral adaptation has focused on the Indo-West Pacific region (Kleypas et al., 2016; Matz et al., 2018, 2020a), corals exist around the world, including throughout the Pacific and in the Caribbean (Veron, 1995). Comparisons among reef systems are important for understanding which results are general and which are sensitive to particular geographic contexts. Here, we implemented a dynamic eco-evolutionary metacommunity model for years 1870-2300 on three spatially realistic reef networks: the Caribbean, Southwest Pacific (SWP) and Coral Triangle (CT). To simulate the response of a coral community to climate change, we modeled the dynamics of two competing coral types with contrasting life-history strategies (Baskett et al., 2014; Darling et al., 2012; Walsworth et al., 2019). In addition to important regional differences, we find that across all three regions, reefs with higher destination strength (larger numbers of immigrating larvae) and lower relative ocean temperatures were most likely to adapt successfully to future warming and that these metrics outperformed other potential predictors of reef persistence.

## Materials and Methods

### General overview

We simulated the cover of two coral types through time during a historical period (1870-2018) and two future sea surface temperature (SST) projections (2018-2300) that followed either RCP 4.5, an emissions reduction and climate stabilization scenario, or RCP 8.5, a scenario without emissions reductions and with continuous and greater warming (Pachauri et al., 2015). Within each region, coral subpopulations exchanged larvae based on previously published biophysical model outputs (Schill et al., 2015; Thompson et al., 2018; Treml et al., 2008) and evolved in response to changing temperatures. To explore the role of ecological vs. evolutionary dynamics on regional and local coral populations, we simulated metapopulation dynamics with different levels of standing genetic variation, which sets evolutionary potential (Norberg et al., 2012; Walsworth et al., 2019). Finally, we constructed general linear models to interpret the influence of patch-level metrics on the minimum coral cover experienced on a reef. Doing so facilitated the identification of temperature and connectivity factors that constrain reef persistence under changing environmental conditions.

### Eco-evolutionary model

We applied an eco-evolutionary model forced with temperature projections and larval connectivity patterns in the Caribbean, SWP and CT coral reef regions. Because reefs are typically considered as distinct habitat patches, we incorporated a novel extension to a continuous-space metapopulation dynamics framework to allow for immigration during the larval phase according to a connectivity matrix **D** that quantifies dispersal connections in each region (described below). The connectivity matrix was based on output from biophysical simulations of coral larval dispersal. We simulated a coral reef metacommunity with two competing coral types: a fast-growing, temperature-intolerant species (‘fast coral’) and a slow-growing, temperature-tolerant species (‘slow coral’) (Darling et al., 2012). Furthermore, growth and mortality rates of each species were affected by the difference between the experienced ocean temperature and an evolving trait called the optimal growth temperature.

On each reef patch *a*, change in coral cover and mean optimal growth temperature were given by

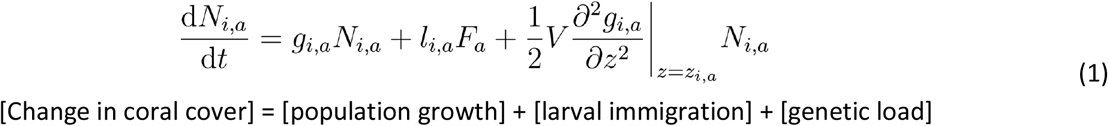

[Change in coral cover] = [population growth] + [larval immigration] + [genetic load]

and

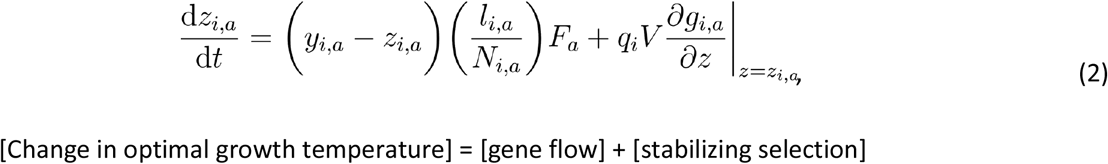

[Change in optimal growth temperature] = [gene flow] + [stabilizing selection]

where *N*_*i,a*_ was the proportion of cover of coral species *i* at site *a* and *z*_*i,a*_ was the optimum growth temperature. The change in coral cover (Eq. 1) for each species was affected by the local population’s growth rate, *g*_*i,a*_, the proportion of free space on the patch, *F*_*a*_, the larval input rate, *l*_*i,a*_, and the additive genetic variance, *V*. Higher *V* makes rapid evolution possible (Lande, 1976) but also leads to a greater genetic load (Kirkpatrick & Barton, 1997). Due to the concavity of the fitness curve with respect to the average trait value and local temperature, genetic load was either negative or zero. The change in optimum growth temperature (Eq. 2) was also a function of the mean population-weighted trait value of immigrants (Hanski et al., 2011), *y*_*i,a*_, and of *q*_*i*_, which reduced the effect of selection at very low coral cover (Norberg et al., 2012). The resident trait was subtracted from the mean trait of the immigrants and scaled by the fraction of cover represented by newly-settled larvae, *l*_*i,a*_ */ N*_*i,a*_, and by free space. Therefore, incoming larvae exerted a stronger effect on the average trait value if they represented a large fraction relative to the current cover of the species and if there was free space for settlement.

In this model, the local population growth rate (Eq. 3) was determined by the intrinsic growth rate, *r*_*i,a*_, mortality rate, *m*_*i,a*_, and competitive interactions encoded in the species interaction matrix α, where α_ij_ was the competitive effect of species *j* on species *i*. In our case, competitive interactions between the fast and slow coral were symmetric such that each species exerted the same effect on the other and intraspecific competition was stronger than interspecific competition (Table S1).

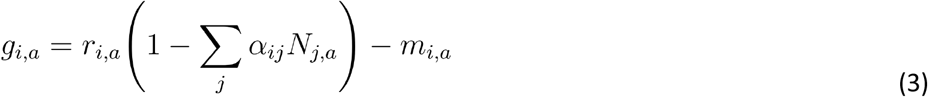

Intrinsic growth (Eq. 4) and mortality (Eq. 5) were Gaussian and exponential functions, respectively, of the local temperature *T*_*a*_, the local average trait value, *z*_*i,a*_, the width of thermal tolerance, *w*_*i*_, and a growth scaling factor, *r*_*0,i*_. Following Walsworth et al. (2019) and the skewed shape of many thermal performance curves (Deutsch et al., 2008), we imposed additional mortality when the current local temperature exceeded the optimum growth temperature, *T*_*a*_ > *z*_*i,a*_ (Eq. 5).

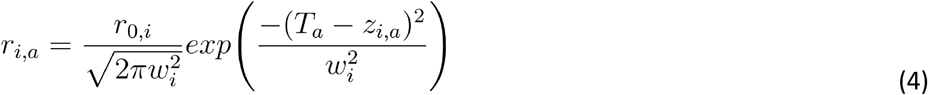

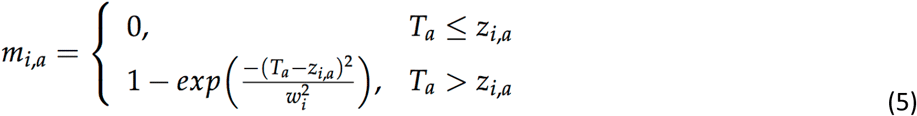

In addition to population dynamics, change in coral cover through time was affected by the dispersal of coral larvae across the network. Our model differed from previous frameworks (Norberg et al., 2012; Walsworth et al., 2019) because we incorporated spatially explicit dispersal among patches instead of a diffusion approximation. Overall, there was higher settlement when there was more free space (i.e., *F*_*a*_ >> 0, see Eq. 1) and a patch without free space (*F*_*a*_=0) had a settlement rate of zero. Free space was the portion of the patch not covered by coral (Eq. 6).

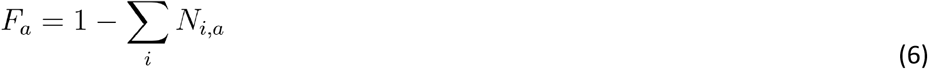

Larval input (*l*_*i,a*_) was calculated from the effective fecundity rate *β* and from the connectivity matrix **D** in which element D_ab_ was the probability of reaching patch *a* from patch *b*. We also accounted for differences in area (*A*_*a*_) among patches such that a small fraction of the reproductive output from a large patch had a disproportionately large effect on a small receiving patch (assuming that there was free space in the latter):

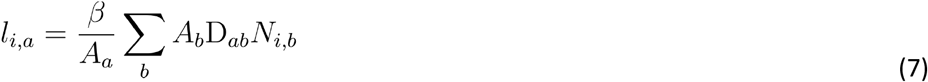

In Eq. 2, the change in mean optimal growth temperature was governed in part by gene flow (first term). The mean incoming trait value for each species (*y*_*i,a*_) was calculated as:

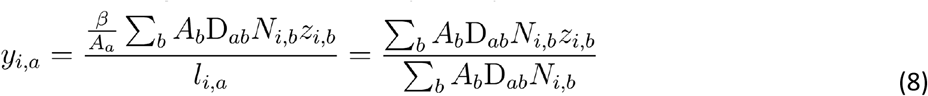

Finally, stabilizing selection,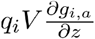, in Eq. 2 acted to match the optimum growth temperature to the local temperature. This evolutionary potential was stronger with increasing *V*, and at very low cover, (*N*_*min*_=10^−6^), *q*_*i*_ reduced the effect of selection (Eq. 9).

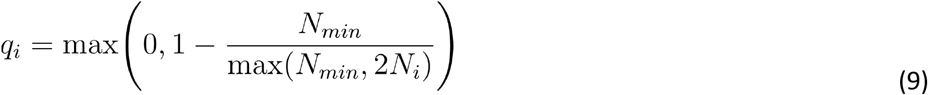

A summary of parameter definitions and values used for simulations is presented in Table S1.

### Potential connectivity

Each region contained many spatially discrete reef patches (423 in Caribbean; 583 in SWP; 2083 in CT), as defined by three unique larval exchange connectivity matrices obtained from previously published biophysical simulations (Caribbean, Schill et al., 2015; Coral Triangle, Thompson et al., 2018; Southwest Pacific, Treml et al., 2008). All connectivity matrices were generated based on simulated larval dispersal for a coral-like species. However, there were differences in the biological assumptions that underlay the creation of each matrix, including a longer pelagic larval duration in the SWP and the absence of larval mortality in the CT (see Table S2 for more details). Although beyond the scope of this study, differences in these key assumptions may have affected the final matrix output. For example, a longer pelagic duration and the lack of larval mortality will tend to increase the number of rare long distance dispersal events.

### Sea surface temperatures

To simulate historical and future ocean warming, spatially explicit SST trajectories were obtained by applying the delta method (Fowler et al., 2007; Hay et al., 2000; Ramirez-Villegas & Jarvis, 2010) to statistically downscale coarser (1×1 degree latitude and longitude) reconstructions from HadISST1 SST for years 1870-2018 (Rayner, 2003) and projections from GISS E2 H for years 2018-2300 (Schmidt et al., 2014) with a climatology created from the higher resolution (0.25×0.25 degree) historical NASA OISST V2 from 1982-2010 (Reynolds et al., 2007). We created SST trajectories for each region under RCP 4.5, an emissions reduction and climate stabilization scenario and RCP 8.5, a scenario without emissions reductions and with continuous and greater warming. Overall, the SST trajectories began in 1870 and ran to 2300, where the reconstruction and projection periods were 1870-2018 and 2018-2300, respectively. Each reef patch experienced a unique thermal environment based on the changing temperature at their location within a grid cell in these trajectories (Fig. S1).

### Simulations

Parameters (Table S1) were chosen to allow coexistence of the fast and slow coral species (Tekwa et al., 2020), which corresponds to empirical observations across the globe (Darling et al., 2012). To facilitate comparison within and among the three regions, all reefs had equivalent parameter values except for area, potential connectivity, and SST time series. To impose a trade-off between the two corals, we set a relatively high growth rate and a narrow thermal tolerance for the fast coral, while the slow coral had a lower growth rate and a wider thermal tolerance (Baskett et al., 2014; Darling et al., 2012). At the beginning of the hindcast run (1870), reefs were initialized such that each coral species started with a cover of 0.25 at every patch (Fig. S2). To examine the effect of evolution on coral persistence, we tested three different levels of additive genetic variance to approximate zero (*V*=0), low (*V*=0.01) and high (*V*=0.1) evolutionary potential. The analyses focused on simulations with an effective fecundity (*β*) of 0.5, although we also calculated trajectories for *β*=0 and *β*=0.05 as a sensitivity test (see Supporting Information) (Álvarez-Noriega et al., 2016).

### Model summaries

We sought to understand whether relatively simple connectivity and temperature metrics could help identify patches that maintained high vs. low coral cover. For each region, patch-level explanatory variables included the change in SST over the 2018-2300 projection period (ΔSST), average SST from 2008-2018 (iSST), the probability of self-connection relative to outgoing connections (local retention, LR), the probability of self-connection relative to incoming connections (self-recruitment, SR), the sum of all incoming connections (destination strength, DS), the difference in mean SST (2008-2018) between the source patches and destination patch of incoming larvae (initial temperature mismatch, ITM), the proportion of incoming connections from patches that are at least 0.5 °C warmer (pr05), and patch area (see Supporting Information for equations). We present a summary of hypothesized ecological and evolutionary effects of each metric in Table 1.

**Table 1.**
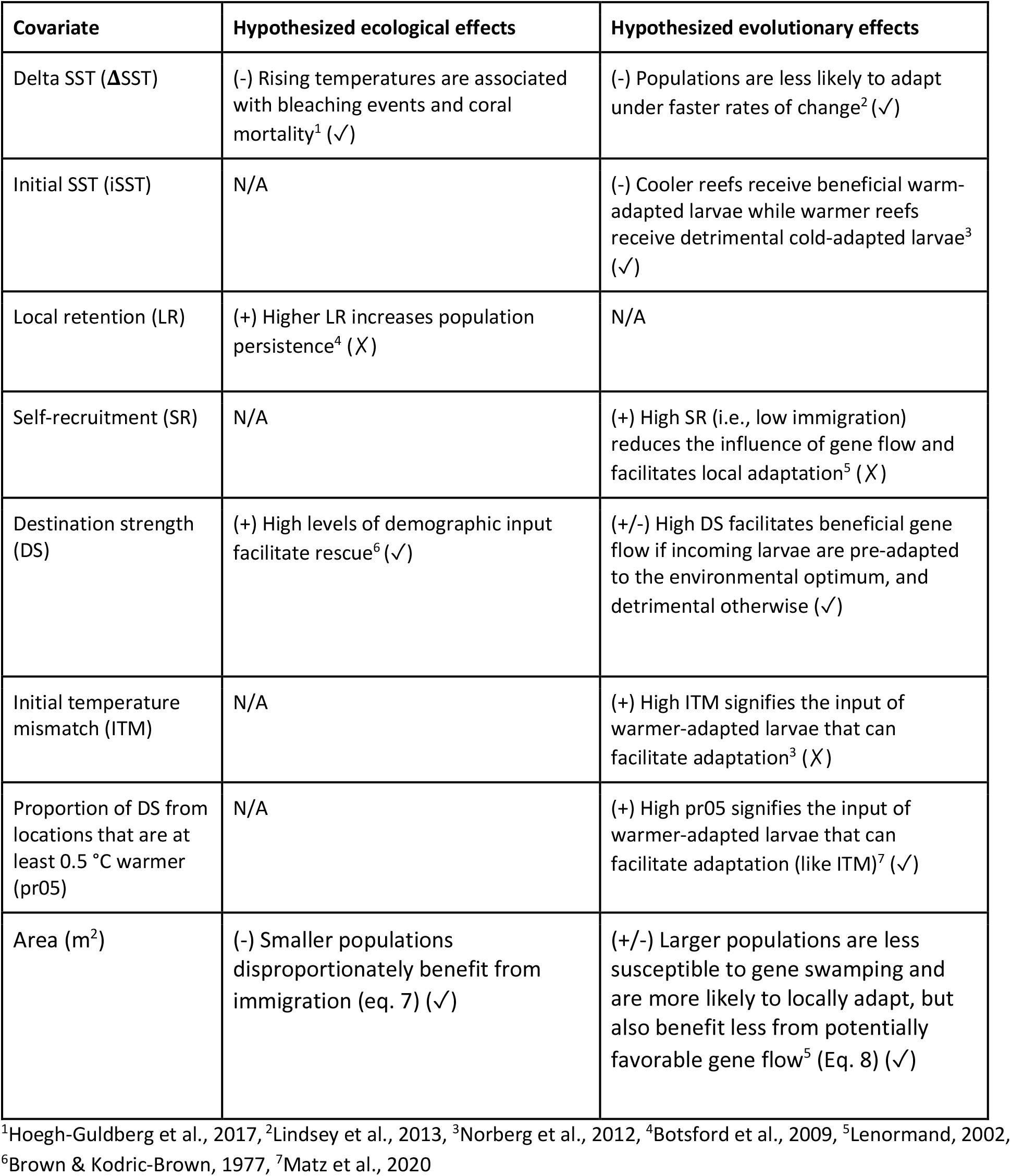
Description of hypothesized ecological and evolutionary effects of site characteristics on coral cover. (+) and (-) indicate a positive or negative effect on coral cover, respectively. (✓) and (✗) indicate that the hypothesis was supported or unsupported, respectively.

| Covariate | Hypothesized ecological effects | Hypothesized evolutionary effects |
| --- | --- | --- |
| Delta SST (ΔSST) | (-) Rising temperatures are associated with bleaching events and coral mortality<sup>1</sup> (✓) | (-) Populations are less likely to adapt under faster rates of change<sup>2</sup> (✓) |
| Initial SST (iSST) | N/A | (-) Cooler reefs receive beneficial warm-adapted larvae while warmer reefs receive detrimental cold-adapted larvae<sup>3</sup> (✓) |
| Local retention (LR) | (+) Higher LR increases population persistence<sup>4</sup> (X) | N/A |
| Self-recruitment (SR) | N/A | (+) High SR (i.e., low immigration) reduces the influence of gene flow and facilitates local adaptation<sup>5</sup> (X) |
| Destination strength (DS) | (+) High levels of demographic input facilitate rescue<sup>6</sup> (✓) | (+/-) High DS facilitates beneficial gene flow if incoming larvae are pre-adapted to the environmental optimum, and detrimental otherwise (✓) |
| Initial temperature mismatch (ITM) | N/A | (+) High ITM signifies the input of warmer-adapted larvae that can facilitate adaptation<sup>3</sup> (X) |
| Proportion of DS from locations that are at least 0.5 °C warmer (pr05) | N/A | (+) High pr05 signifies the input of warmer-adapted larvae that can facilitate adaptation (like ITM)<sup>7</sup> (✓) |
| Area (m<sup>2</sup>) | (-) Smaller populations disproportionately benefit from immigration (eq. 7) (✓) | (+/-) Larger populations are less susceptible to gene swamping and are more likely to locally adapt, but also benefit less from potentially favorable gene flow<sup>5</sup> (Eq. 8) (✓) |

<sup>1</sup>Hoegh-Guldberg et al., 2017, <sup>2</sup>Lindsey et al., 2013, <sup>3</sup>Norberg et al., 2012, <sup>4</sup>Botsford et al., 2009, <sup>5</sup>Lenormand, 2002,
<sup>6</sup>Brown & Kodric-Brown, 1977, <sup>7</sup>Matz et al., 2020

To quantify the relative association of patch-level characteristics with coral cover, we fit generalized linear models with binomial errors to the minimum fractional coral cover of each patch in a region between 2018-2300 (Burnham et al., 2002; Zuur et al., 2009). We applied a log transformation to LR, SR, DS, pr05, and area (log(x+0.001)), and a log-modulus transformation for ITM (sign(x) * log(|x|+1)) to minimize skew. We centered and scaled all variables (mean=0, SD=1) after transformations. In addition to exploring the full statistical model, the model average, and the model with the lowest Akaike information criterion (AIC; Burnham & Anderson, 2004), we created statistical models for three hypotheses: 1) Connectivity-only with LR, SR, and DS; 2) Temperature-only with ΔSST and iSST; and 3) Warm-adapted gene flow with pr05 and DS. Inferences regarding the influence of each of these variables on coral cover were made based on the coefficients of each explanatory variable in the full statistical model. To help avoid issues with applying statistical methods to simulated data (White et al., 2014), we limited the interpretation of these results to a qualitative comparison of effect size among our metrics and did not conduct statistical significance testing.

## Results

### Future Coral Cover

Across all regions, we found that evolution was critical in maintaining coral cover through warming. Corals persisted under both mild (RCP 4.5) and severe (RCP 8.5) warming scenarios with high genetic variance (*V*=0.1) (Fig. 1a-c, Table S3). With intermediate genetic variance (*V*=0.01), corals managed to persist under mild warming (RCP 4.5) in the Caribbean and CT but not under strong warming (RCP 8.5), while in the SWP, corals maintained a small amount of cover even under severe warming (RCP 8.5) (Fig. S3). For moderate genetic variance (*V*=0.01), warming was projected to cause coral declines until 2050, followed by slow recovery that did not reach historical levels by 2300 (Fig. 1). In general, higher genetic variance (*V*) led to higher minimum and final coral cover across each region (Fig. 1). The Coral Triangle retained the highest coral cover, while the Caribbean had the lowest cover across all *V*.

**Figure 1.**
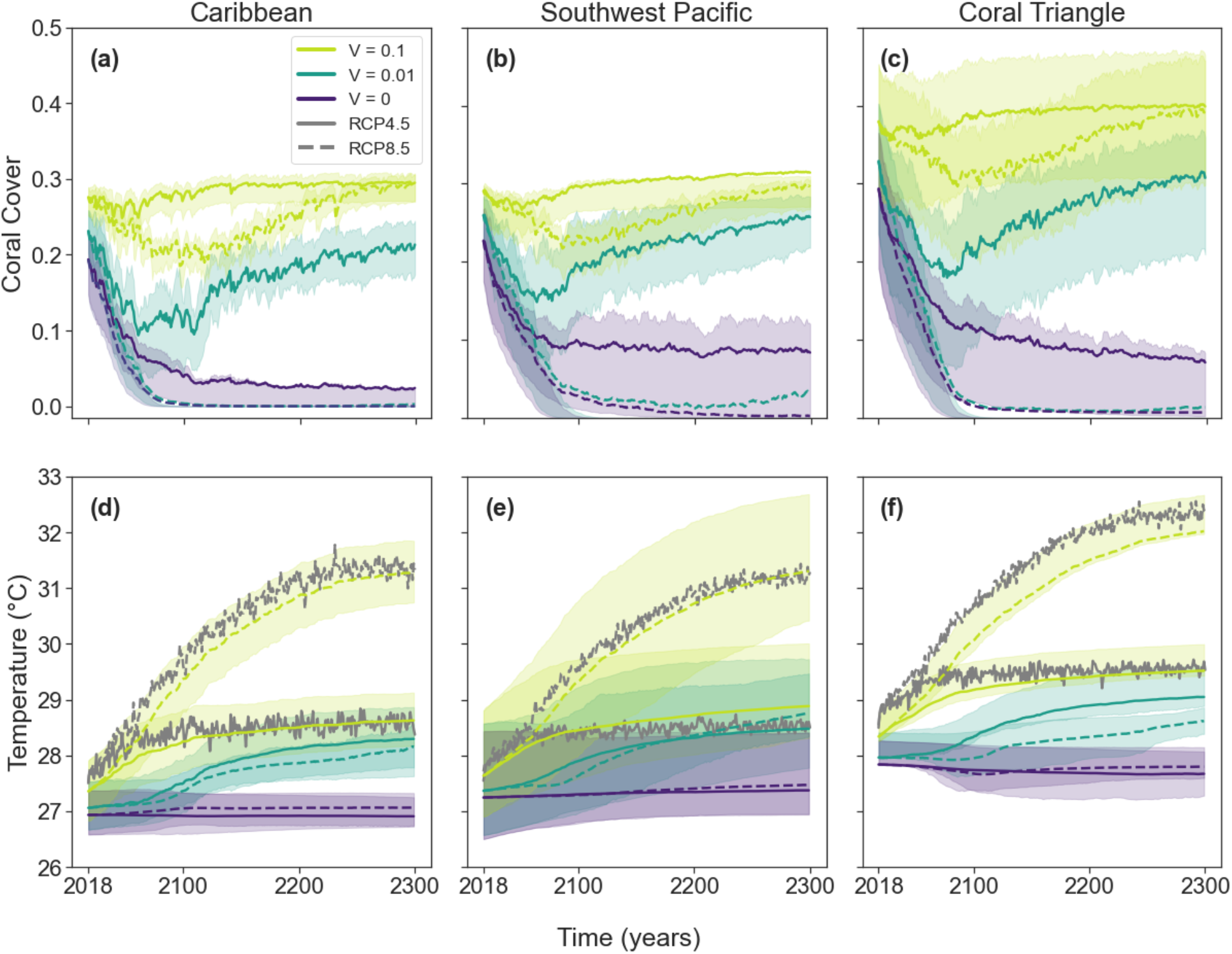
Mean total coral cover (sum of both functional types) (a-c) and mean trait values with sea surface temperatures (d-f) averaged across all reefs in the Caribbean, Southwest Pacific and Coral Triangle under RCP 4.5 (solid lines) and RCP 8.5 (dashed lines). Trajectories are shown at three levels of additive genetic variance: *V*=0 (violet), *V*=0.01 (blue-green) and *V*=0.1 (yellow-green). Error bounds refer to the 25th and 75th percentiles among reefs. Mean SSTs across each network are shown in gray (d-f).

Across regions, substantial spatial variation in coral cover was apparent under mild warming (Fig. 2, Fig. S4, Fig. S5). In the Caribbean, the sites with the highest coral cover were found near the Bahamas and south of Cuba. In the Southwest Pacific, the high cover reefs were near American Samoa and the area between Papua New Guinea and the Solomon Islands. In the Coral Triangle, the highest cover reefs were near the Paracel Islands in the South China Sea and the Greater Sunda Islands in Indonesia. However, spatial patterns were different by the end of the more severe warming scenario (RCP 8.5) in the Caribbean and the Coral Triangle (Fig. S4). In the Caribbean, only Bermuda and two sites off the coast of South America had surviving corals, while in the Coral Triangle, only sites near Hainan, China and in the southern Great Barrier Reef maintained coral cover by the end of the projection (Fig. S4 b,d,f). Sensitivity testing without larval fecundity or with lower fecundity (*β*=0 or *β*=0.05, respectively) revealed that higher fecundity (in effect, more larval dispersal) increased the variation of minimum coral cover values across each network (Fig. S6).

**Figure 2.**
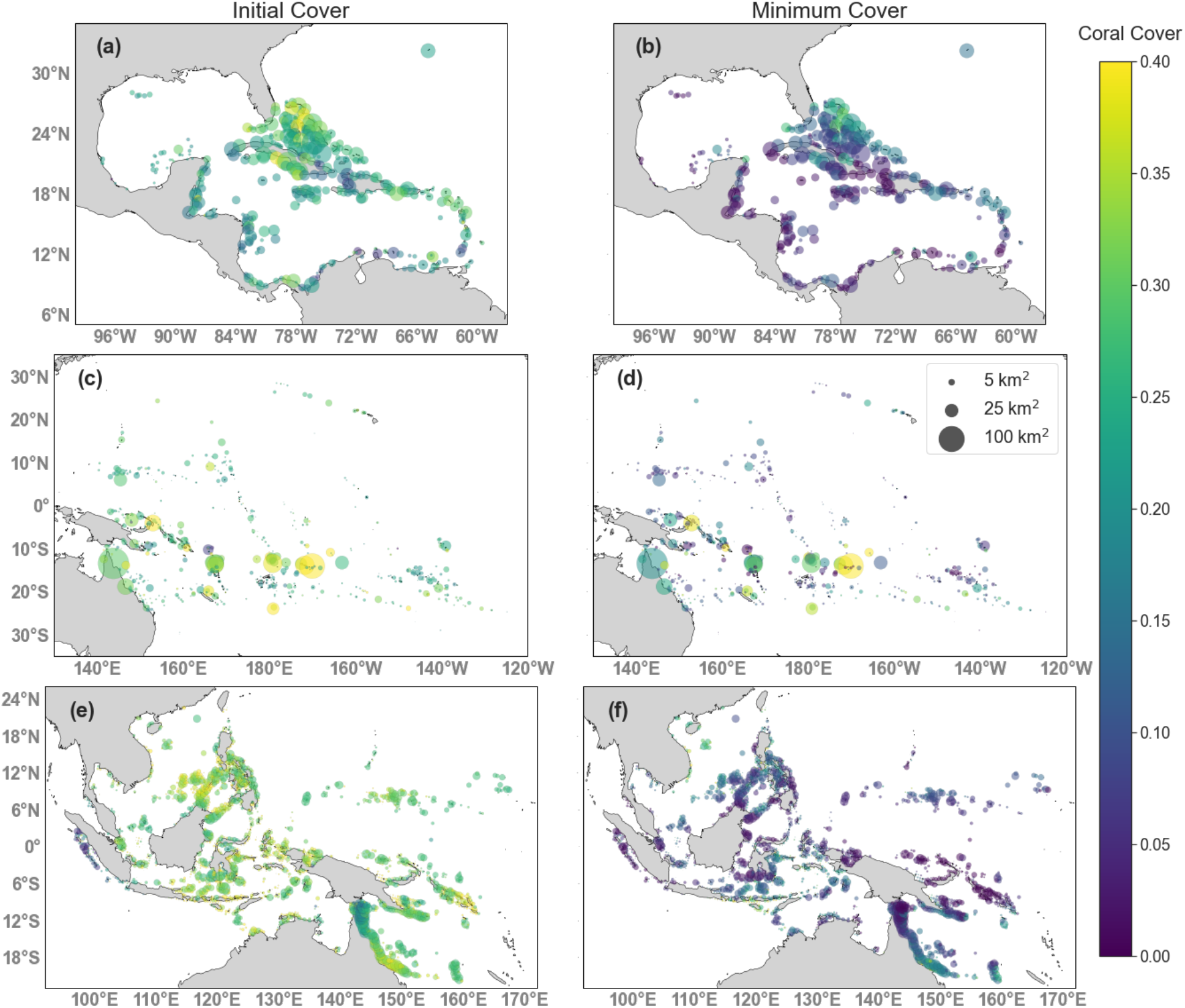
Coral cover in 2018 (a, c, e) and minimum cover under RCP 4.5 (b, d, f) for the Caribbean (a, b), Southwest Pacific (c, d), and Coral Triangle (e, f). Reef sizes are proportional to areas. Results are shown for *V*=0.01 and *β*=0.5.

We also found a strong temporal correlation in coral cover across all regions such that reefs with higher cover in 2018 tended to also have a higher minimum cover and a higher final cover. We found that initial and final coral cover was highly correlated with coefficients of 0.89, 0.95, and 0.85 for the Caribbean, SWP, and CT, respectively (Fig. S7 a). Initial and minimum cover (Fig. S7 b) and final and minimum cover (Fig. S7 c) were similarly highly correlated.

The simulations predicted time-dependent shifts in coral composition in response to warming. Slow-growing, temperature-tolerant corals were relatively more abundant during the initial projection period and when coral cover was lowest across regions (Fig. 3 a-c). In contrast, fast-growing corals with narrower thermal tolerance recovered faster and were more abundant later in the simulation period (Fig. 3a-c, Table S3). Slow-growing, stress-tolerant corals had more variable cover among reefs for all regions throughout the time series, even when they were less abundant than fast-growing corals (Table S3). Fast-growing corals were able to adapt to local temperatures by 2300 across regions, whereas slow-growing corals continued to have optimal temperatures that were lower than experienced temperatures (Fig. 3d-f). In other words, the mismatch between optimal trait and temperature were better tolerated initially by slow-growing corals, but higher evolutionary rates in fast-growing corals were ultimately advantageous because they were able to match their optimal trait values to the local environment.

**Figure 3.**
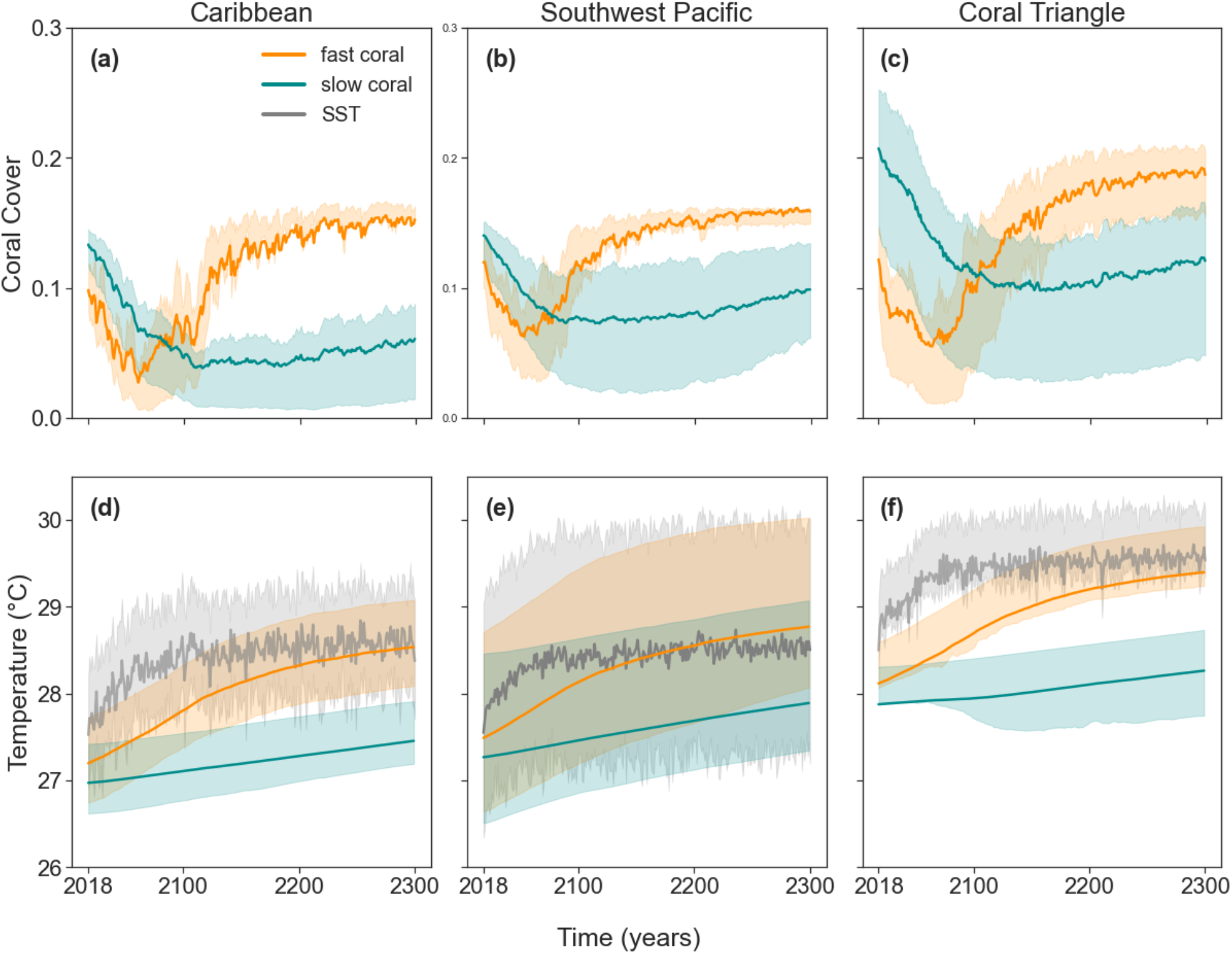
Projected coral cover through time under RCP 4.5 by species (fast = orange; slow = cyan) for the (a) Caribbean, (b) Southwest Pacific, and (c) Coral Triangle. (d)-(f) Mean sea surface temperature (gray) and trait value for each species. Additive genetic variance V=0.01. Error bounds show the 25th and 75th percentiles among reefs.

### Predictors of Patch-scale Coral Cover

The strongest predictor of minimum coral cover across regions and levels of additive genetic variance was an individual reef’s destination strength (DS) (Fig. 4a). This network metric, which corresponds to the sum of a reef’s incoming dispersal links, was positively correlated with minimum cover across all regions, levels of *V*, and warming scenarios (Fig. 4, Fig. S8, Fig. S9). Minimum coral cover was also negatively associated with the initial sea surface temperature (iSST) across all regions, levels of *V*, and warming scenarios (Fig. S8, Fig. S9).This latter pattern was in agreement with our evolutionary hypothesis that initially hot reefs generally received maladapted (colder adapted) larvae from connected reefs and were less able to adapt to warming temperatures (Table 1).

**Figure 4.**
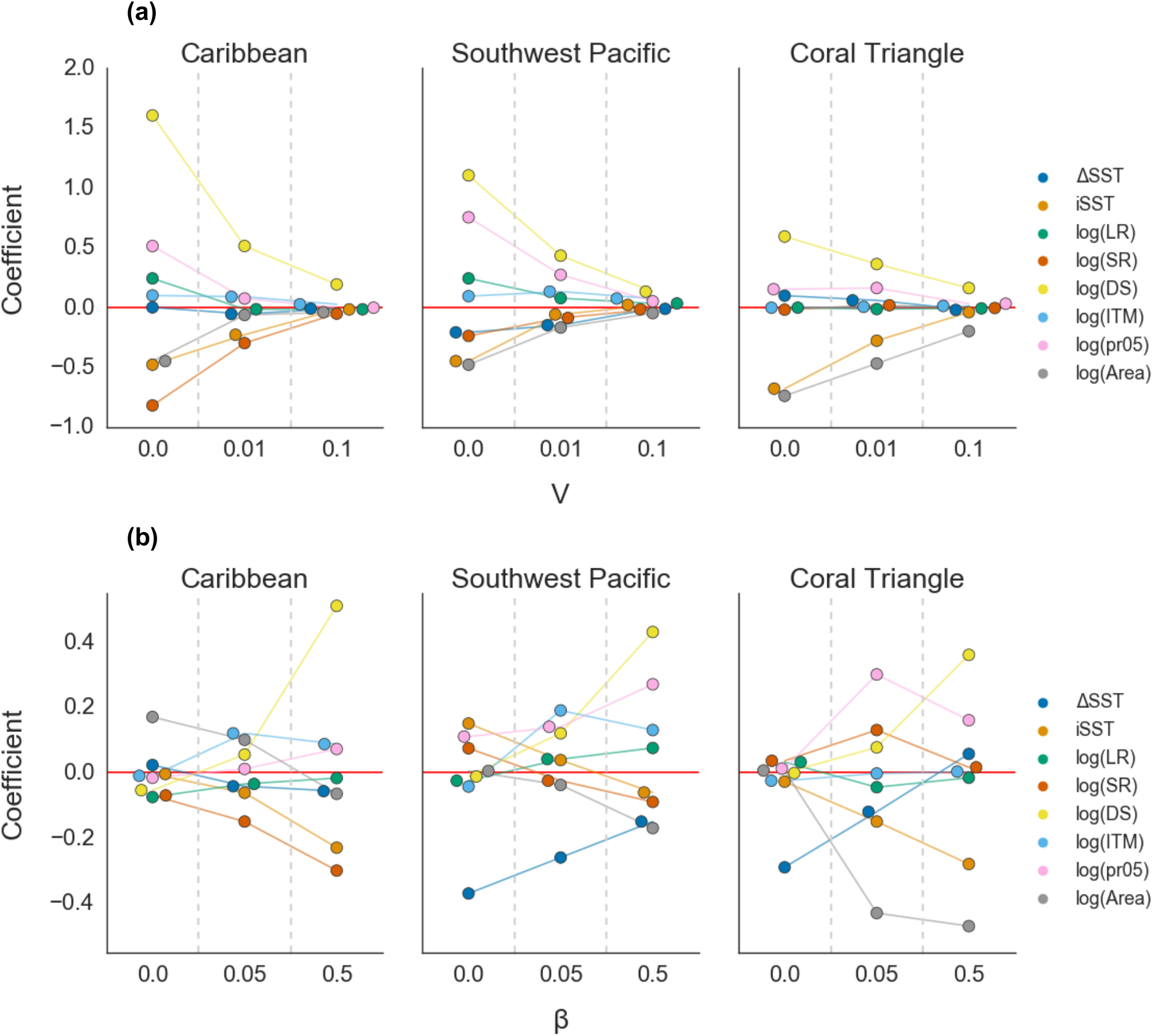
Standardized coefficient (effect size) of each covariate on minimum coral cover (a) across levels of genetic variance or (b) levels of fecundity in the Caribbean, Southwest Pacific and Coral Triangle. ΔSST is the change in sea surface temperature over the projection period, iSST is the initial temperature at the start of the projection, LR is local retention, SR is self-recruitment, DS is destination strength, ITM is initial temperature mismatch and pr05 is the proportion of incoming links from sites that are at least 0.5 °C warmer (based on iSST). Results are for RCP 4.5 and *β*=0.5 in part (a) and for *V*=0.01 for part (b).

Certain predictors had a strong association with minimum cover in one region, but not in the others. For example, pr05 was strongly positively correlated with minimum coral cover in the SWP, moderately correlated in the Caribbean only with no additive genetic variance, and only weakly positively associated in the CT (Fig. 4). In the Caribbean, self-recruitment (SR) was strongly negatively correlated with minimum cover but was only weakly negatively correlated in the SWP and had a near-zero effect size in the CT (Fig. 4). Across all metrics, the effect sizes for all covariates decreased as *V* increased (Fig. 4).

Some predictors differed in their frequency distributions across regions (Fig. S10). For example, the Caribbean had higher local retention of larvae than the CT and SWP. The SWP had a wider range of ΔSST and iSST than the other two regions, and subsequently, a higher proportion of cold sites. The CT had substantially higher DS, and a greater number of rare, extremely small sites.

As a sensitivity test, we also performed simulations with no effective fecundity (*β*=0) which meant that there was no dispersal among reefs. In this scenario, only within-reef ecological and evolutionary processes were important. Unsurprisingly, connectivity metrics had weak and inconsistent associations with minimum cover across regions without dispersal (Fig. 4b). In addition, the negative iSST effect disappeared at low fecundity, supporting mechanistic hypotheses that the iSST effect was mediated by dispersal of larvae among sites, rather than being a result of within-site dynamics (Table 1).

By comparing alternate statistical models based on three hypotheses (1) *Connectivity-only* with LR, SR, and DS; 2) *Temperature-only* with ΔSST and iSST; and 3) *Warm-adapted gene flow* with pr05 and DS), we found that the *Warm-adapted gene flow* model performed better than the *Temperature-only* and *Connectivity-only* models across all regions based on △AIC and R^2^ values (Table S4, Table S5, Table S6).

## Discussion

Reefs around the world are projected to experience frequent severe bleaching and mortality events during this century (Donner, 2009; Frieler et al., 2013; Logan et al., 2014; van Hooidonk et al., 2013). Our model provides further evidence of corals’ capacity to evolutionarily adapt and recover within 200 years under lower greenhouse gas emissions scenarios, but that such recovery would not be possible without evolution (Bay et al., 2017; Matz et al., 2020; Walsworth et al. 2019). Coral populations in our model collapsed with no additive genetic variance (*V*=0), and we found that the declines were less severe under more rapid evolution. This finding held across three major coral reef networks with different dispersal patterns and temperature trajectories. In addition, we found that an initial loss of fast-growing corals over the coming century may be offset by their faster recovery if thermal conditions stabilize. Lastly, we found that differences in minimum coral cover were associated with both larval connectivity and temperature, including the quantity of incoming larvae and relative SST, which can help inform conservation strategies designed to maintain coral cover.

Several regional differences in population dynamics were apparent in our simulations. For example, there was higher initial coral cover, slower recovery, and more evolutionary lag (a larger mismatch between coral traits and local temperature) in the Coral Triangle, which has a larger number of reefs and greater rates of potential larval recruitment. The lowest minimum coral cover occurred in the Caribbean, which had fewer reefs and high local retention of larvae. The Caribbean also experienced the longest pre-recovery period. The Southwest Pacific had the least evolutionary lag and correspondingly recovered to near-historical levels of coral cover by 2300. The SWP experienced the least warming on average, had the greatest variation in temperature and warming across the network, and had a higher proportion of cool sites, all of which may have contributed to the region’s recovery.

Previous studies also projected spatial variation in coral declines. Couce et al. (2013) implemented a statistical habitat suitability model to project global coral habitat suitability in response to ocean warming and acidification for the years 2010, 2040, and 2070. They found the greatest declines in the Western Pacific Warm Pool, which corresponds to much of the Coral Triangle and northern SWP in our model where we also found marked declines in cover. The regions projected to maintain high suitability for corals in Couce et al.’s model also correspond to most of the regions which maintain cover in our model (e.g., Southern Great Barrier Reef in both the CT and SWP regions, Greater Sunda Islands, South China Sea in the CT, American Samoa and the Solomon Islands in the SWP). However, the Couce et al. model projects that the entire Caribbean will maintain high suitability, or even increase in suitability, while our model projects severe declines in cover throughout much of the region. Differences in our results are expected due to the imperfect correlation between habitat suitability and abundance, as the former ignores all biological processes, as well as the coarser resolution of their model inputs (1×1 degrees). Using an individual-based model based on forward genetic simulations, Matz et al. (2020) tracked coral metapopulations under warming in the Indo West-Pacific. This study projected a similar spatial distribution of declines across the Coral Triangle as seen in our model and the Couce et al. results, with the highest declines occurring in near equatorial reefs and higher maintenance of coral cover in northwestern and southeastern reefs away from the equator. However, Matz et al. projected less severe coral cover declines in both warming scenarios as compared to our model. This could be attributed to differences in the way that reproduction, dispersal and genetic variation were specified in our two models, as well as the combination of parameter values used in the simulations.

To our knowledge, no other study has modelled multiple regional coral networks under an eco-evolutionary framework, and thus the regional differences suggested by our model stand to be tested. The observed regional differences in population dynamics are likely due to intrinsic differences in temperatures and dispersal among the regional networks, but may also be due, in part, to methodological differences in the models used to generate the connectivity relationships in our model (Schill et al., 2015; Thompson et al., 2018; Treml et al., 2008). These results suggest that future research and management should consider the unique characteristics of each region and that there is a need for additional regional comparison studies. While difficult to implement, consistent dispersal simulation approaches across regions would assist with these comparisons.

Our results suggest that shifts in coral community composition in response to increasing temperatures should be expected and may be reversed during coral population recovery. In our simulations, the fast-growing coral with a narrow temperature tolerance experienced greater initial declines but also exhibited a higher capacity for adaptation relative to the slow-growing species. Our fast-growing coral closely resembles branching corals from the family Acroporidae (Darling et al., 2012). While this may imply that acroporid populations in the Caribbean can eventually recover when warming stabilizes, we note that the observed declines in real populations are primarily attributed to disease (Aronson & Precht, 2001), herbivore die-offs (Lessios, 2016) and local stressors (Cramer et al., 2020), all of which we did not model, in addition to thermal stress (Hughes, 1994). We also do not model the possibility of range expansion in response to increases in temperature, which has been observed in the acroporid fossil record (Baird et al., 2012; Precht & Aronson, 2004; Yamano et al., 2011). Additionally, genomic analyses indicate that acroporids experienced a period of population decline following the Mid-Pleistocene Transition (global cooling), and then a period of rapid diversification and population growth following the Northern Hemisphere Glaciation (global warming and sea level rise) (Mao et al., 2018). While our model indicates that acroporids may be more sensitive to short-term (years to decades) shifts in temperature than other scleractinian families, their fossil record and genome indicate that they may be poised for long-term range expansion in a warmer climate, given their survival.

We examined whether temperature and larval connectivity characteristics were useful for explaining which reefs would persist with high cover and which would not. Analyses of past bleaching events have found that temperature-based metrics, including mean SST, temporal variability of SST, and degree heating weeks (a measure of thermal stress) are strong predictors of past coral bleaching and mortality at specific sites (Hughes et al., 2018; McClanahan & Maina, 2003; Safaie et al., 2018; Sully et al., 2019; Welle et al., 2017). Studies that link larval connectivity patterns to marine population persistence typically focus on the network scale and calculate centrality metrics to identify sites which disproportionately contribute to metapopulation growth (Kininmonth et al., 2019; Treml & Halpin, 2012; Watson et al., 2011). Our work focused on integrating these two types of metrics with additional site-specific connectivity measures such as destination strength (Thompson et al., 2018), self-recruitment, and local retention (Burgess et al., 2014; Hastings & Botsford, 2006), as several previous studies have asserted that larval settlement and recruitment rates directly limit the local population size (Caley et al., 1996; Terence P Hughes, 1990; Menge et al., 2003). This comprehensive assessment not only highlighted the independent contributions of temperature and connectivity to coral persistence and adaptation, but also the importance of their interactions for future reef persistence when there is potential for an evolutionary response.

Our results suggest that, while evolution has a broadly positive effect on coral persistence at all sites and across all regions, variation exists among sites’ resilience to ocean warming due to differences in temperature and connectivity characteristics. We found a consistently positive effect of a reef’s destination strength and a consistently negative effect of initial SST on minimum cover during warming, regardless of region or additive genetic variance. Recently, Matz et al. (2020) found that pre-warming SST and the fraction of recruits immigrating from sites that were at least 0.5°C warmer (pr05) were strong predictors of reef persistence in the Indo-West Pacific. Larvae that were pre-adapted to warmer conditions strongly benefited cooler reefs, consistent with genetic theory (Norberg et al., 2012). In line with these results, we found that initial SST and pr05 were relatively effective predictors in our model, although neither was as effective as destination strength. The positive effect of destination strength likely operated through the ecological effects of larval immigration, implying that the demographic benefits of connectivity outweighed the potential negative evolutionary effects of gene swamping (Lenormand, 2002). Furthermore, the negative effect of initial SST was likely due to warmer reefs receiving cold-adapted larvae: as the network warmed, cold-adapted larvae arriving in relatively warm reefs counteracted evolutionary adaptation (Norberg et al., 2012). These interactions of ecological and evolutionary processes highlight the importance of considering both to understand future reef states.

Though destination strength and initial SST had consistent effects across all regions, we also found that the types of reefs most likely to survive future warming differed among regions in important ways. For example, self-recruitment had a stronger negative association with cover in the Caribbean, and area had a stronger negative effect in the Coral Triangle. Self-recruitment is an indication of the ‘openness’ of a reef, or the amount of larval input from the rest of the network relative to the contribution from the reef itself. In other words, reefs with high self-recruitment received fewer larvae from locations other than their own. The Caribbean’s prevailing surface currents tend to move larvae from warm sites to cold sites or other warm sites, but rarely from cold to warm (Carrillo et al., 2015; Chollett et al., 2012). Thus, open reefs in the Caribbean generally received evolutionarily neutral or beneficial larvae and only rarely received maladapted larvae. Relatively closed reefs in this region did not experience the synergistic benefits of demographic support and assisted evolution from warm-adapted larvae. Next, the stronger negative association of area with minimum cover in the Coral Triangle was likely due to the wider range of reef areas in this region, including the presence of extremely small sites that were much less abundant in other regions. Therefore, this region was more likely to contain connections between sites with a large difference in area. Because the quantity of larvae dispersing in our model scaled with both reef area and coral cover, larger sites were more likely to demographically support smaller sites than vice versa, leading to a strong association between reef size and minimum cover in the CT. Again, these results suggest that future research and management would benefit from considering the unique characteristics of each region.

Evolution mitigated the impact of the environmental variation among sites in each regional network. As additive genetic variance decreased, the magnitudes of coefficients associated with all network factors increased. In other words, with reduced evolutionary capacity, the local temperature and connectivity of each site had a greater effect on its coral cover. In contrast, all sites tended to have higher cover if evolutionary capacity was high. Thus, the maintenance of genetic variation in coral networks would help support coral persistence across a range of environments. With little evolutionary potential, we can expect that well-connected small reefs in colder microclimates are most likely to persist under warming.

Our projections contained a number of important assumptions. We assumed that within-reef coral dynamics were identical among reefs and regions, except for temperature and network connectivity. This assumption allowed us to investigate evolutionary, network, and environmental effects on coral cover, holding other characteristics constant. In reality, coral communities in different regions are composed of different species (beyond our model of fast and slow corals) that have varied responses to environmental stress (Darling et al., 2012). We also assumed that the corals’ thermal optima and growth rate were not correlated to other traits which affect fitness; however, the rate of adaptation seen in our model may not be possible if thermal tolerance trades off with other traits affecting fitness (Etterson & Shaw, 2001). In addition, we only tracked coral cover in two dimensions, which ignored three-dimensional reef structural complexity that plays a prominent role in ecosystem services (Darling et al., 2019). Some coral communities may also exhibit ecological alternative stable states when interactions with macroalgae are included (Mumby et al., 2007). In this study, we did not include non-coral species and chose to parameterize the model to ensure coexistence, rather than alternative stable states between the corals (Tekwa et al., 2020). While this choice allowed for outcomes that were not as sensitive to initial conditions, one interesting avenue for future investigation would be the interaction of evolution with ecological alternative stable states. Eco-evolutionary feedbacks have been linked to alternative community compositions in lake systems (Strauss, 2014; Walsh et al., 2012) and may have similar impacts on coral reefs (Mumby et al., 2007). Finally, the timescale of initial decline in coral cover and subsequent recovery can only be interpreted qualitatively. Our results suggest that eventual recovery is possible with evolution, but it is not possible to infer from our model when recovery will occur. That is because recovery timescales are affected by both growth rates and additive genetic variance, which are difficult to measure and may vary greatly across species and regions (but see Anderson et al., 2017; Carilli et al., 2010; D’Croz & Maté, 2004; Dziedzic et al., 2019; Edmunds, 2005 for estimates of coral growth rates; Jokiel & Coles, 1977; Kirk et al., 2018 for estimates of additive genetic variance). Lastly, we note that our results are limited to assessing the response of individual reefs to warming; the impact of any particular reef on the network is beyond the scope of this work.

Our results suggest that some general strategies are likely beneficial for coral conservation under warming. First, limiting greenhouse gas emissions and hence warming will facilitate coral adaptation, as demonstrated by the stark differences in coral cover between RCP4.5 and RCP8.5. Second, evolutionary potential is critical for mitigating coral loss and facilitating recovery of corals around the world, both during and after warming. Thus, policies that maintain genetic diversity are likely to have important long-term benefits. For example, implementing protection across environmentally distinct sites can maintain relatively high additive genetic variance at the network scale by preserving populations that are locally adapted to different thermal regimes (Baums et al., 2019; Howells et al., 2013). Because higher genetic variance also leads to persistence at local scales, genotyping approaches to quantify local genetic variation can help inform conservation and restoration efforts (e.g., protecting sites with high diversity) (Baums et al., 2019). Third, our results identify the characteristics of reefs that are likely to maintain coral cover vs. those that will likely experience significant coral declines. For example, relatively cool reefs with high larval input may have a greater chance of coral cover maintenance or recovery in response to conservation measures that aim to mitigate external stressors such as reductions in local sedimentation (Bégin et al., 2016; Dubinsky & Stambler, 1996) and nutrient input (Dubinsky & Stambler, 1996). On the other hand, warm reefs with low larval input may benefit the most from larval supplementation (Cruz & Harrison, 2017) or restoration efforts (Baums et al., 2019; Ladd et al., 2018) since they are predicted to have less recovery potential overall. This finding also has implications for reef managers in terms of site selection criteria for management interventions. Managers could intentionally aim to include some cooler reefs and those with high larval settlement (resistant reefs), as well as some hotter reefs and those with low larval settlement (vulnerable reefs) within the managed network. Including a portfolio of reef types within a managed network helps to facilitate multiple means of adaptation to warming, including evolutionary and demographic rescue, in addition to local adaptation (Mumby et al., 2011; Walsworth et al., 2019).

Our projections suggest that a future for corals is possible if warming is limited. Maintaining evolutionary potential and habitat connectivity are both important for the continued existence of coral populations. While we predict a sharp decline in reef cover, we also expect recovery with sufficient genetic variability under a less severe warming scenario. In this work, we linked individual reef characteristics to coral cover response in three major reef networks. Future work can build on these results to investigate how conservation strategies could harness adaptive potential across the reef network as a whole under climate change.

## Acknowledgements

We gratefully acknowledge funding from the Gordon and Betty Moore Foundation and The Nature Conservancy. We also thank Lukas DeFilippo and members of the Pinsky lab for helpful discussions on earlier versions of this manuscript.

