## Supporting Information for "Evolution and connectivity influence the persistence and recovery of coral reefs under climate change in the Caribbean, Southwest Pacific, and Coral Triangle"

**Supporting Methods**

*Sensitivity analysis for effective fecundity ($\beta$)*

Due to high uncertainty and variability surrounding the effective fecundity of corals (Álvarez-Noriega et al., 2016), we calculated trajectories for $\beta$=0 and $\beta$=0.05 in addition to the default $\beta$=0.5 (Fig. S6). In the main text, we focus on $\beta$=0.5 because this sets the contribution of larval input to coral growth to be approximately two orders of magnitude less than the contribution of clonal growth. At equilibrium and assuming all patches are equal (in temperature, area, and connections), clonal growth contributes approximately $\frac{r_0}{\sqrt{2\pi w^2}}$, which is the maximum growth rate (Table S1). Larval settlement contributes approximately mean(DS)*$\beta$ to overall population growth, where DS is the destination strength of a patch. For the fast and slow coral, the maximum growth rates (i.e., maximum clonal growth rates) are 0.6 and 0.2, respectively. From the connectivity matrices, mean(DS) values for the Caribbean, Southwest Pacific and the Coral Triangle were 0.12, 0.08 and 0.28, respectively. Using $\beta$=0.05 or 0.5, the larval contribution to population growth in the Caribbean would be 0.006 and 0.06 for the two levels, 0.004 and 0.04 in the Southwest Pacific and 0.014 and 0.14 in the Coral Triangle. As such, the clonal growth contribution to coral cover increase is roughly one to two orders of magnitude greater than the contribution from larval establishment in the Caribbean and Southwest Paciifc. In the Coral Triangle, clonal growth is the same order of magnitude or one order greater than larval settlement.

*Equations to compute site characteristics*

For the following calculations, **D** is the connectivity matrix where element $D_{ab}$ is the probability of reaching site *a* from site *b*. $A_a$ is the area of patch *a* and $SST_{a,t}$ is the temperature experienced at site *a* during year *t*.

Delta SST (**ΔSST**):

$$\Delta\mathrm{SST}_a = \frac{1}{10}\sum_{t=2290}^{2299}\mathrm{SST}_{a,t} - \frac{1}{10}\sum_{t=2008}^{2017}\mathrm{SST}_{a,t}$$

Initial SST (**iSST**):

$$\mathrm{iSST}_a = \frac{1}{10}\sum_{t=2008}^{2017}\mathrm{SST}_{a,t}$$

Local retention (**LR**) was the probability of self-connection relative to the sum of all outgoing connections:

$$LR_a = \frac{\mathrm{D}_{aa}}{\sum_b \mathrm{D}_{ba}}$$

Self-recruitment (**SR**) was the probability of self-connection relative to the sum of all incoming connections:

$$SR_a = \frac{\mathrm{D}_{aa}}{\sum_b \mathrm{D}_{ab}}$$

Destination Strength (**DS**) was the sum of all incoming connections (including self-connection):

$$DS_a = \sum_b \mathrm{D}_{ab}$$

Initial temperature mismatch (**ITM**) was the average mismatch between temperatures in a focal reef and its source reefs, scaled by connection strength and area, averaged over the final 10 years of the hindcast (2008-2017). Here, *n* is the total number of patches in the network.

$$ITM_a = \frac{1}{10nA_a} \sum_{t=2008}^{2018} \sum_b (\mathrm{SST}_{b,t} - \mathrm{SST}_{a,t}) \mathrm{D}_{ab} A_b$$

**pr05** was the proportion of DS from locations that were at least 0.5 °C warmer (Matz et al. 2020). The difference in 10-year mean SSTs between source site *b* and the destination site *a* is $\delta_{ab}$, and $DS_{05,a}$ is the sum of incoming connections to site *a* from all sites *b* such that $\delta_{ab} \geq 0.5$°C.

$$\delta_{ab} = \frac{1}{10} \sum_{t=2008}^{2018} SST_{b,t} - \frac{1}{10} \sum_{t=2008}^{2018} SST_{a,t}$$

$$DS_{05,a} = \sum_b DS_{ab}$$ for all *b* for which $\delta_{ab} \geq 0.5$°C.

$$pr05_a = \frac{DS_{05,a}}{DS_a}$$

**Supporting Tables**

**Table S1.** Parameter definitions and values used in simulations.

| Parameter | Definition | | Fast coral | Slow coral |
|---|---|---|---|---|
| $r_{0,i}$ | Scaling factor for growth rate; Note that maximum growth rate is $\frac{r_0}{\sqrt{2\pi w^2}}$ | | 1.5 | 1.5 |
| $w_i$ | Thermal tolerance breadth | | 1.0 | 3.0 |
| $\beta$ | Effective fecundity | | 0, 0.05, 0.5 | 0, 0.05, 0.5 |
| $V$ | Additive genetic variance | | 0, 0.01, 0.1 | 0, 0.01, 0.1 |
| $\alpha$ | Competition matrix | Fast coral | 5.77 | 0.9 |
| | | Slow coral | 0.9 | 5.77 |

**Table S2**. Methods and parameters used to create the connectivity matrices.

| **Item Description** | **Caribbean** | **Southwest Pacific** | **Coral Triangle** |
|---|---|---|---|
| Publication | (Schill et al., 2015) | (Treml et al., 2008) | (Thompson et al., 2018) |
| Number of Reef Units | 423 | 583 | 2083 |
| Reef location and area determination | Millenium Coral Reef Mapping Project for reef locations and areas, reviewed by in-country reef experts | Digital Chart of the World Server plus Spalding et al. 2001 & Oliver et al. 2004 | Global Distribution of Coral Reefs (UNEP-WCMC), which merges data from the Millennium Coral Reef Mapping Project and the World Atlas of Coral Reefs |
| Ocean Circulation Model | NOAA Real-Time Ocean Forecast System [RTOFS] database (8 sq. km. resolution) (Rivin & Mehra 2010) | NOAA Environmental Modeling Center's ocean analysis system (25 sq. km. resolution) (Ji et al. 1995) | Regional Ocean Modelling System developed for the Coral Triangle region [CT-ROMS] (Castruccio et al. 2013) |
| Maximum Pelagic Larval Duration | 30 days | 60 days | 30 days |
| Larval Mortality | 20% per day | 4% per day | none |
| Spawning events | Two per year from 2008-2011, starting on the last dates of the quarter moon (August - September) | One mass spawning season - September through November 2001 | Biannual mass spawning events lasting 5 days (spring and fall) |
| Pre-competency Period | Gamma cumulative function allowing all larvae to reach full competency in 3 days | none | Beginning at 3 days with full competency at 10 days |
| Settlement behavior | After reaching competency, larvae over coral habitat settled at a rate of 75% per day. | If the density of larvae at a downstream reef site exceeded 1 per cell, a connection was made between the two sites | none |
| Local density & fecundity | Amount of larvae released proportional to reef area | 10,000 larvae per sq. km. | 25 particles from each of the oceanic grid cells within a reef site, for a maximum of 8000 particles released from each site. |

**Table S3**. Summary statistics of coral cover (total, fast coral only, slow coral only) across reefs at beginning and end of the RCP 4.5 projection, as well as the minimum cover each site experienced for $V$ = 0.01 and $\beta$ = 0.5 . Values are shown as the average +/- the standard deviation across sites.

| | **2008-2018** | **Minimum** | **2290-2300** |
|---|---|---|---|
| **Caribbean (both species)** | 0.262 (0.053) | 0.083 (0.064) | 0.209 (0.065) |
| **SWP (both species)** | 0.275 (0.061) | 0.127 (0.099) | 0.256 (0.081) |
| **CT (both species)** | 0.348 (0.126) | 0.150 (0.124) | 0.309 (0.171) |
| **Caribbean (fast coral)** | 0.122 (0.028) | 0.025 (0.025) | 0.151 (0.032) |
| **SWP (fast coral)** | 0.132 (0.031) | 0.048 (0.041) | 0.159 (0.038) |
| **CT (fast coral)** | 0.136 (0.058) | 0.043 (0.052) | 0.189 (0.089) |
| **Caribbean (slow coral)** | 0.140 (0.032) | 0.035 (0.036) | 0.059 (0.047) |
| **SWP (slow coral)** | 0.144 (0.036) | 0.065 (0.059) | 0.097 (0.058) |
| **CT (slow coral)** | 0.212 (0.084) | 0.080 (0.070) | 0.120 (0.105) |

**Table S4**. General linear model summary for the Caribbean region with $V = 0.01$ and $\beta = 0.5$. Relative Variable Importance (RVI) is the sum of model weights over all models including each explanatory variable. Values in parentheses are standard errors.

| | RVI | Full | Best | Warm Larvae | Connectivity | Temperature |
|---|---|---|---|---|---|---|
| **∆SST** | 0.94 | -0.057 (0.41) | -0.055 (0.41) | | | -0.057 (0.39) |
| **iSST** | 1 | -0.23 (0.51) | -0.23 (0.51) | | | -0.25 (0.4) |
| **log(LR)** | 0.28 | -0.017 (0.78) | | | -0.16 (0.76) | |
| **log(SR)** | 1 | -0.3 (1.1) | -0.32 (0.68) | | -0.11 (1.1) | |
| **log(DS)** | 1 | 0.51 (0.86) | 0.52 (0.82) | 0.2 (0.38) | 0.35 (0.71) | |
| **log(ITM)** | 1 | 0.087 (0.33) | 0.088 (0.33) | | | |
| **log(pr05)** | 0.97 | 0.072 (0.5) | 0.072 (0.5) | 0.25 (0.39) | | |
| **log(Area)** | 0.86 | -0.065 (0.54) | -0.063 (0.54) | | | |
| **∆AIC** | | 1.9 | 0 | 200 | 290 | 290 |
| **$R^2$** | | 0.68 | 0.68 | 0.47 | 0.35 | 0.34 |
| **Akaike Weight** | | 0.22 | 0.57 | $1.10 \times 10^{-44}$ | $1.70 \times 10^{-63}$ | $2.60 \times 10^{-64}$ |

**Table S5**. General linear model summary for the Southwest Pacific region with *V* = 0.01 and *β* = 0.5. See legend for Table S4 for further description.

| | RVI | Full | Best | Warm Larvae | Connectivity | Temperature |
|---|---|---|---|---|---|---|
| **∆SST** | 1 | -0.15 (0.36) | -0.15 (0.36) | | | -0.29 (0.32) |
| **iSST** | 1 | -0.061 (0.38) | -0.061 (0.38) | | | -0.22 (0.29) |
| **log(LR)** | 1 | 0.075 (0.48) | 0.075 (0.48) | | 0.099 (0.46) | |
| **log(SR)** | 1 | -0.09 (0.54) | -0.09 (0.54) | | -0.15 (0.53) | |
| **log(DS)** | 1 | 0.43 (0.51) | 0.43 (0.51) | 0.29 (0.33) | 0.43 (0.38) | |
| **log(ITM)** | 1 | 0.13 (0.23) | 0.13 (0.23) | | | |
| **log(pr05)** | 1 | 0.27 (0.39) | 0.27 (0.39) | 0.37 (0.31) | | |
| **log(Area)** | 1 | -0.17 (0.4) | -0.17 (0.4) | | | |
| **∆AIC** | | 0 | 0 | 560 | 1400 | 1200 |
| **$R^2$** | | 0.98 | 0.98 | 0.95 | 0.78 | 0.83 |
| **Akaike Weight** | | 1 | 1 | $2.40x10^{-121}$ | $6.50x10^{-300}$ | $2.60x10^{-262}$ |

**Table S6**. General linear model summary for the Coral Triangle region with *V* = 0.01 and *β* = 0.5. See legend for Table S4 for further description.

| | RVI | Full | Best | Warm Larvae | Connectivity | Temperature |
|---|---|---|---|---|---|---|
| **∆SST** | 1 | 0.057 (0.3) | 0.057 (0.3) | | | 0.083 (0.29) |
| **iSST** | 1 | -0.28 (0.3) | -0.28 (0.3) | | | -0.22 (0.25) |
| **log(LR)** | 0.4 | -0.016 (0.47) | | | 0.0011 (0.44) | |
| **log(SR)** | 0.4 | 0.016 (0.49) | | | 0.058 (0.46) | |
| **log(DS)** | 1 | 0.36 (0.4) | 0.36 (0.35) | 0.29 (0.34) | 0.29 (0.37) | |
| **log(ITM)** | 0.33 | 0.0029 (0.17) | | | | |
| **log(pr05)** | 1 | 0.16 (0.34) | 0.16 (0.33) | 0.27 (0.29) | | |
| **log(Area)** | 1 | -0.47 (0.26) | -0.46 (0.25) | | | |
| **∆AIC** | | 2.7 | 0 | 7600 | 9400 | 9900 |
| **$R^2$** | | 1 | 1 | 0.84 | 0.62 | 0.52 |
| **Akaike Weight** | | 0.072 | 0.28 | 0 | 0 | 0 |

### Supporting Figures

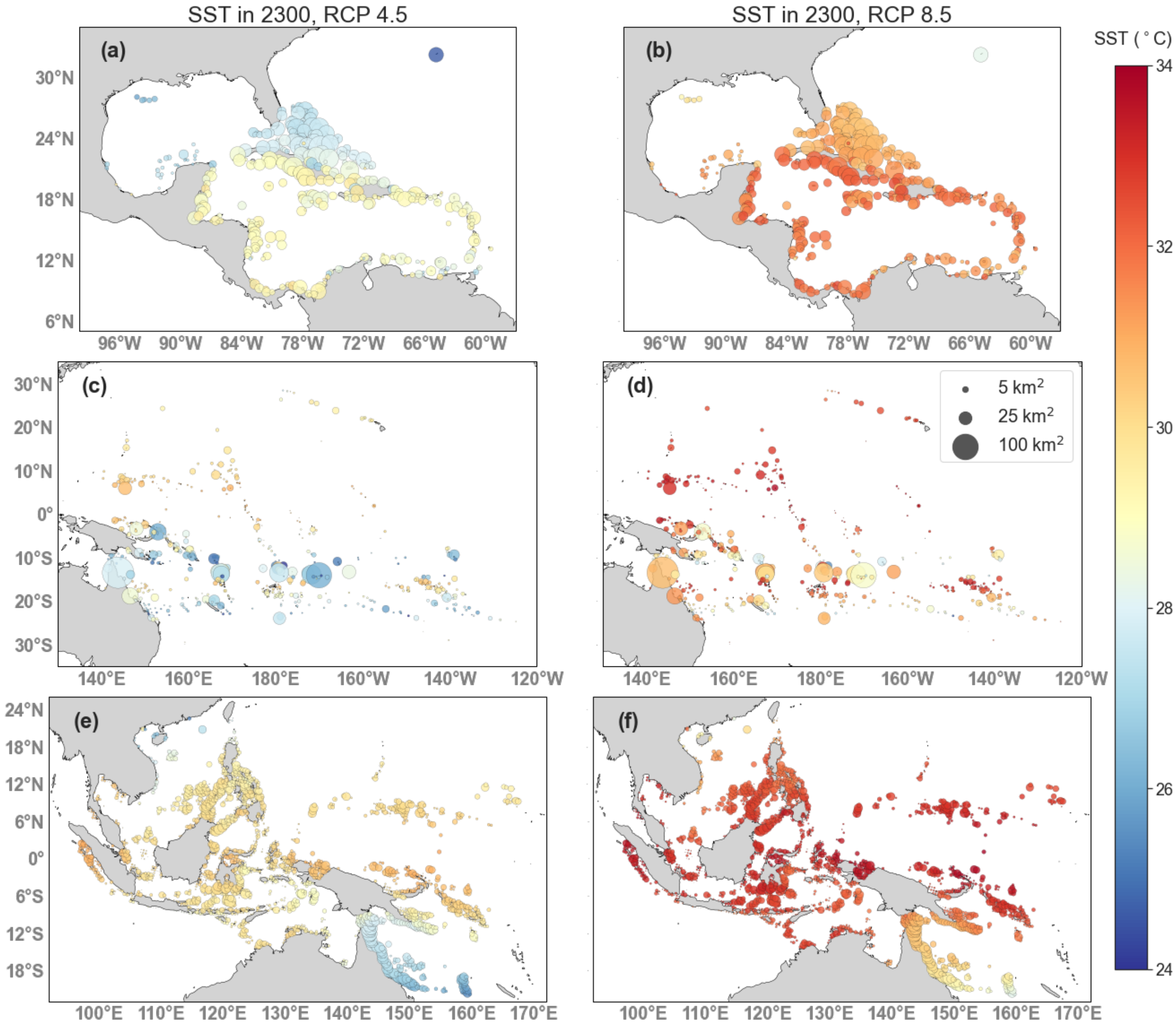


**Figure S1.** Sea surface temperature at the end of the projection period (year 2300) under RCP 4.5 (a,c,e) and RCP 8.5 (b,d,b) for the Caribbean (a,b), Southwest Pacific (c,d, and Coral Triangle (e,f). Dots for reef patches are scaled by reef area (see legend).

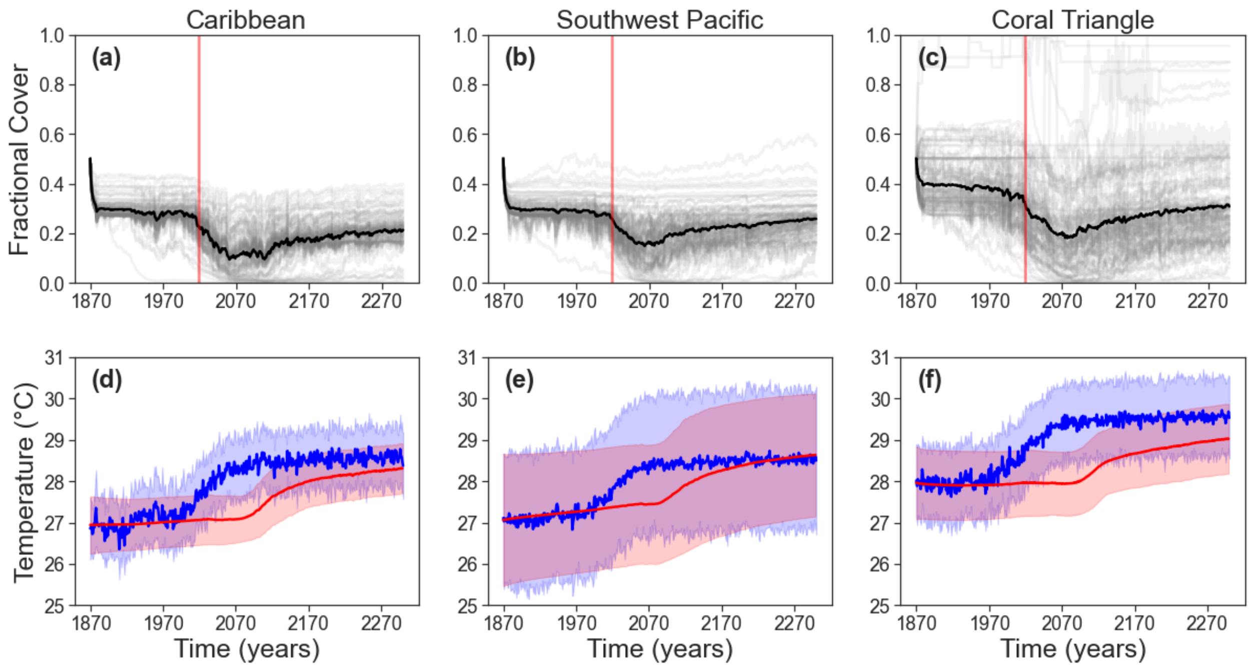


**Figure S2.** Coral cover through time (including hindcast) for years 1870 to 2300 under RCP 4.5 for the (a) Caribbean, (b) Southwest Pacific (b), and (c) Coral Triangle. Dark black lines represent mean trajectories for the whole network, gray lines represent individual reef trajectories for 100 randomly selected sites, and the vertical red line marks the beginning of the projection period. (d)-(f) Network-averaged temperature (blue) and trait value (red). Shaded regions represent one standard deviation above and below the mean. All reefs started at 0.5 cover, with 0.25 cover for each coral type.

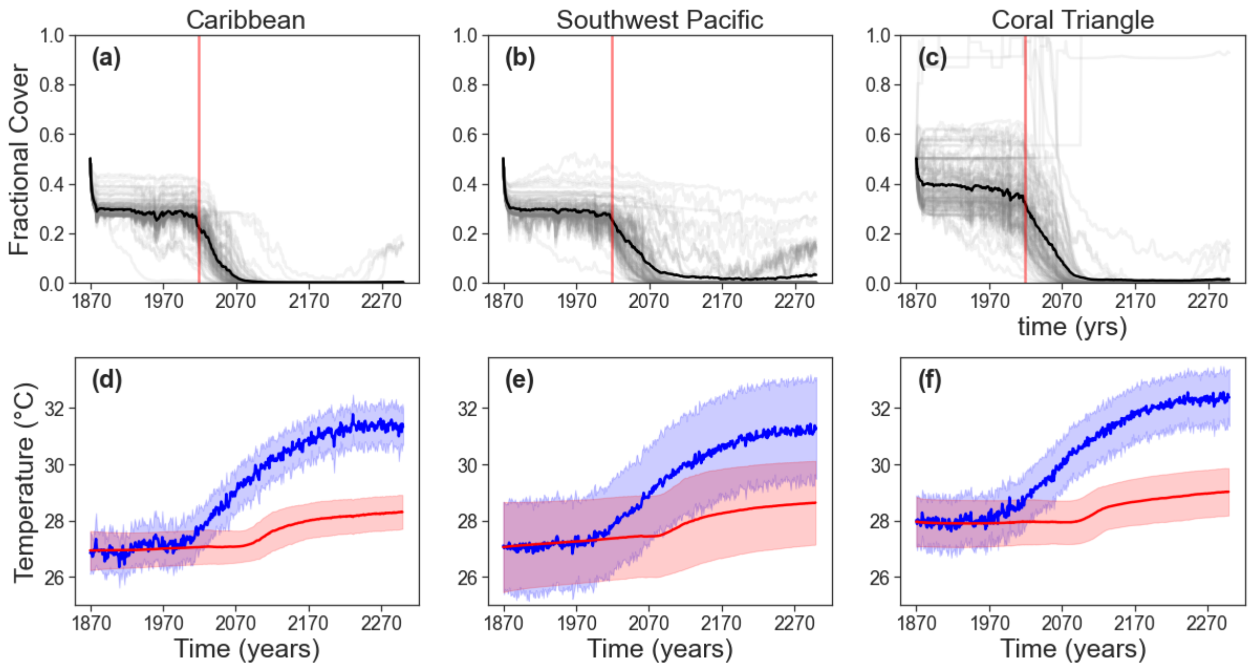


**Figure S3.** Coral cover through time (including hindcast) for years 1870 to 2300 under RCP 8.5 for the (a) Caribbean, (b) Southwest Pacific, and (c) Coral Triangle. Dark black lines represent mean trajectories for the whole network, gray lines represent individual reef trajectories for 100 randomly selected sites, and the vertical red line marks the beginning of the projection period. (d)-(f) Network-averaged temperature (blue) and trait value (red). Shaded regions represent one standard deviation above and below the mean. All reefs started at 0.5 cover, with 0.25 cover for each coral type.

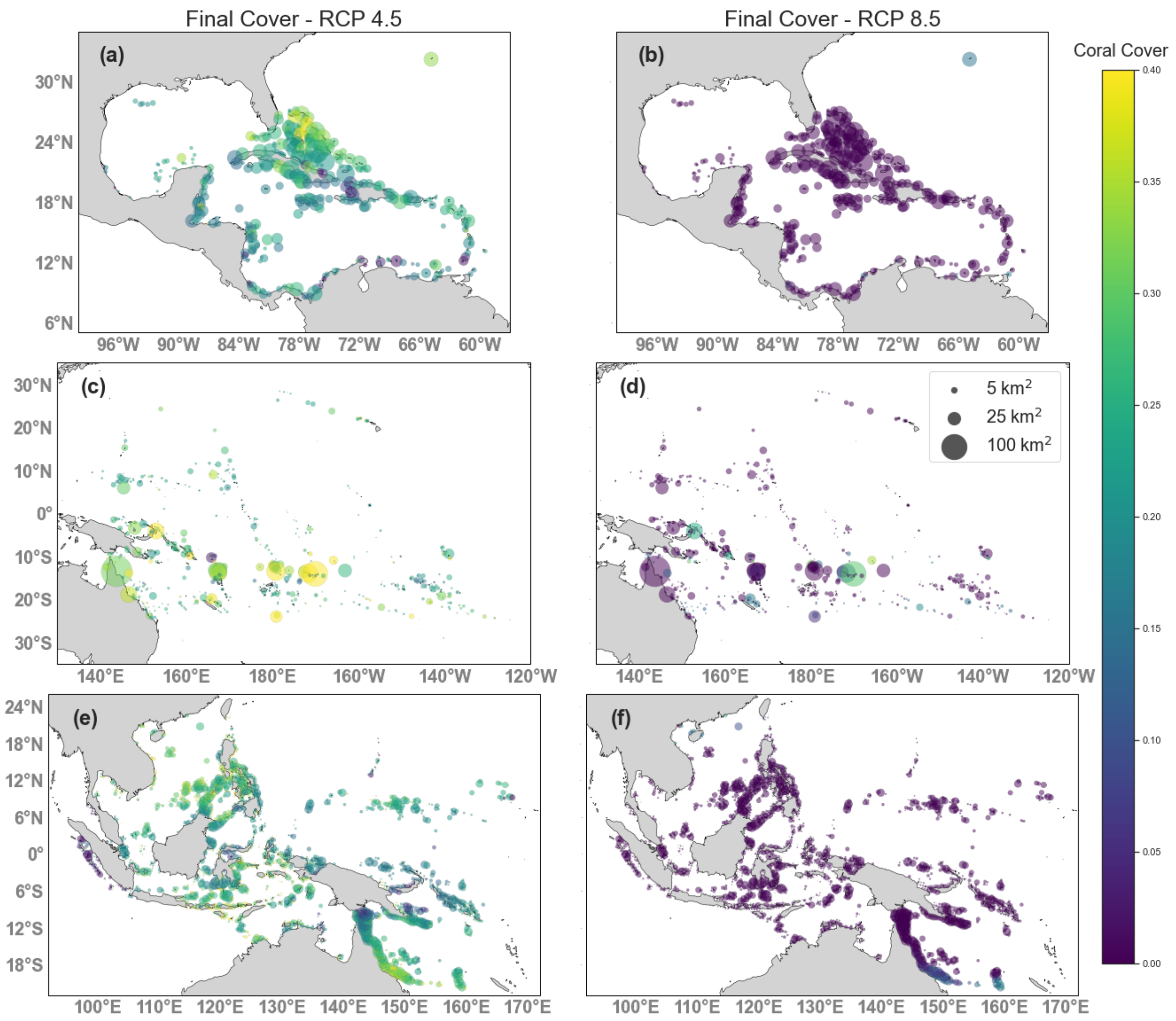


**Figure S4.** Coral cover at the end of the projection period (year 2300) with $V$=0.01 and $\beta$=0.5 under RCP 4.5 (a,c,e) and RCP 8.5 (b,d,b) for the Caribbean (a,b), Southwest Pacific (c,d, and Coral Triangle (e,f).

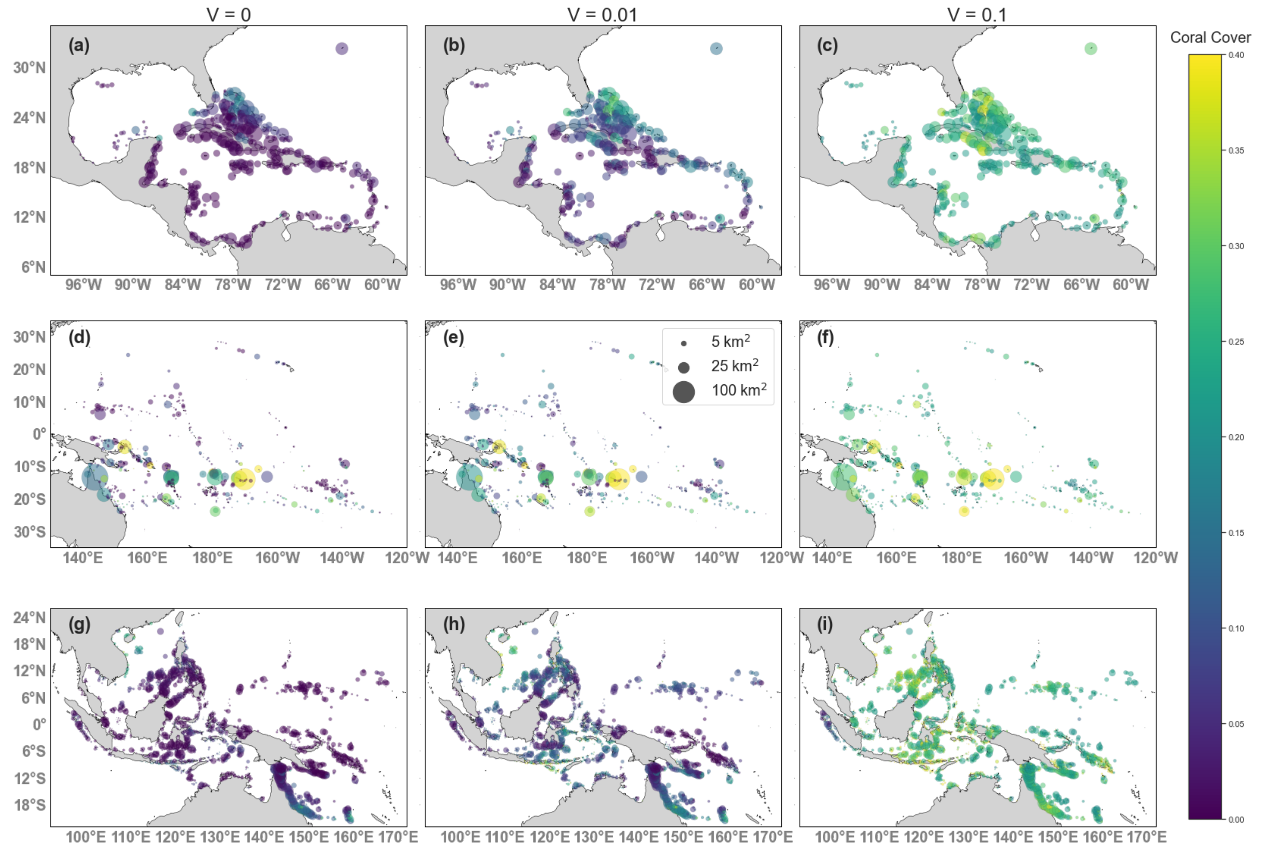


**Figure S5.** Minimum coral cover with three levels of *V* under RCP 4.5 for the Caribbean (a-c), Southwest Pacific (d-f) and Coral Triangle (g-i).

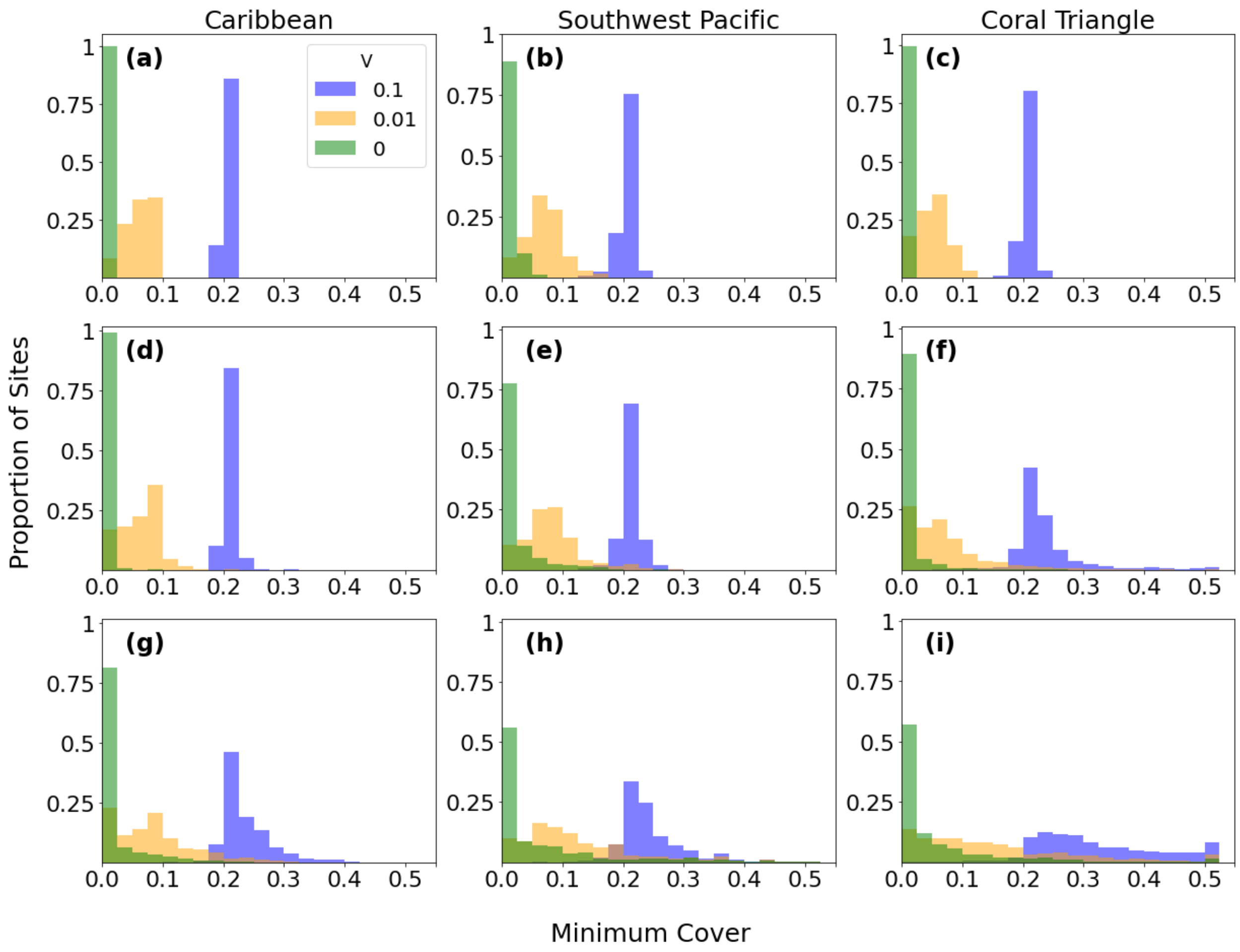


**Figure S6.** Distribution of minimum cover in each region across the three values of additive genetic variance (V) for *β* = 0 (a-c), *β* = 0.05 (d-f), and *β* = 0.5 (g-i) under RCP 4.5.

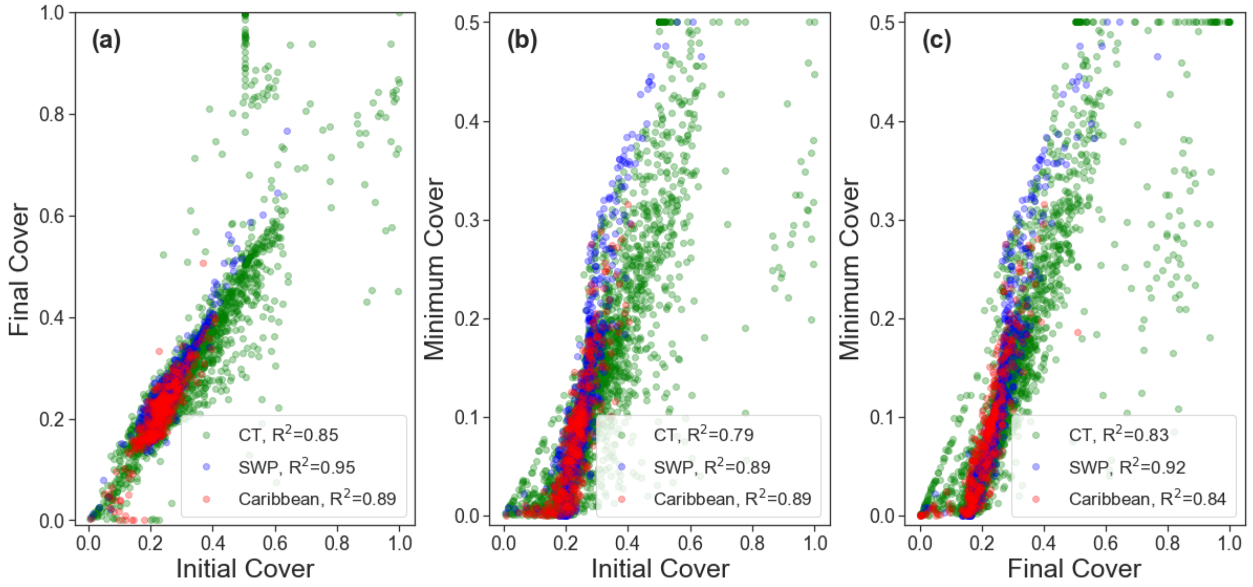


**Figure S7.** Correlation plots of initial (2018) and final (2300) cover (A), initial and minimum cover (B), and final and minimum cover (C) in the Caribbean (red), Southwest Pacific (blue) and Coral Triangle (green).

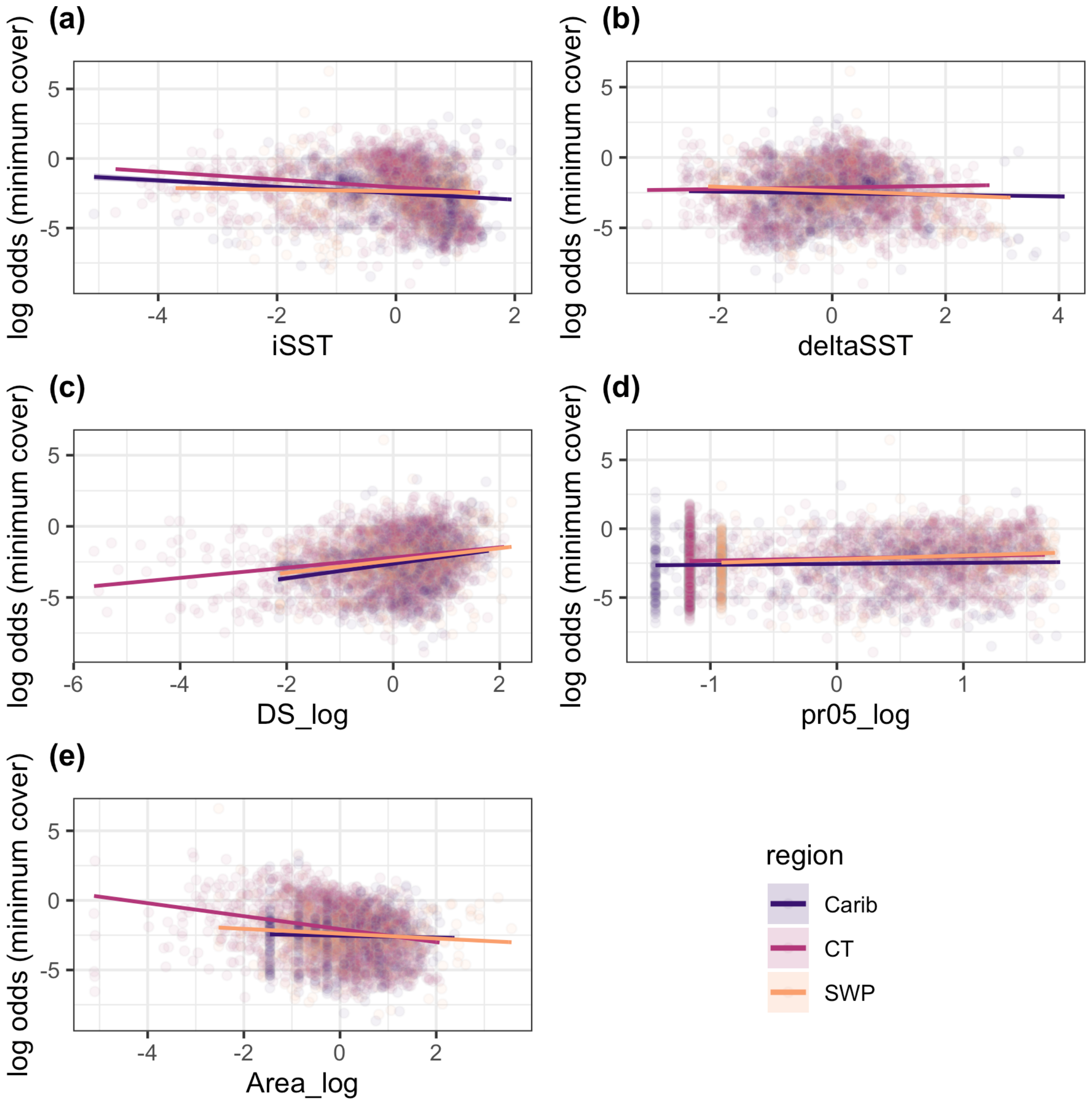


**Figure S8.** Partial effects regression showing the influence of iSST (a), **Δ**SST (b), destination strength (c), initial trait mismatch (d), and area (e) on minimum coral cover for all three regions with V=0.01. Error bars show the 95% confidence intervals but are generally too small to be visible.

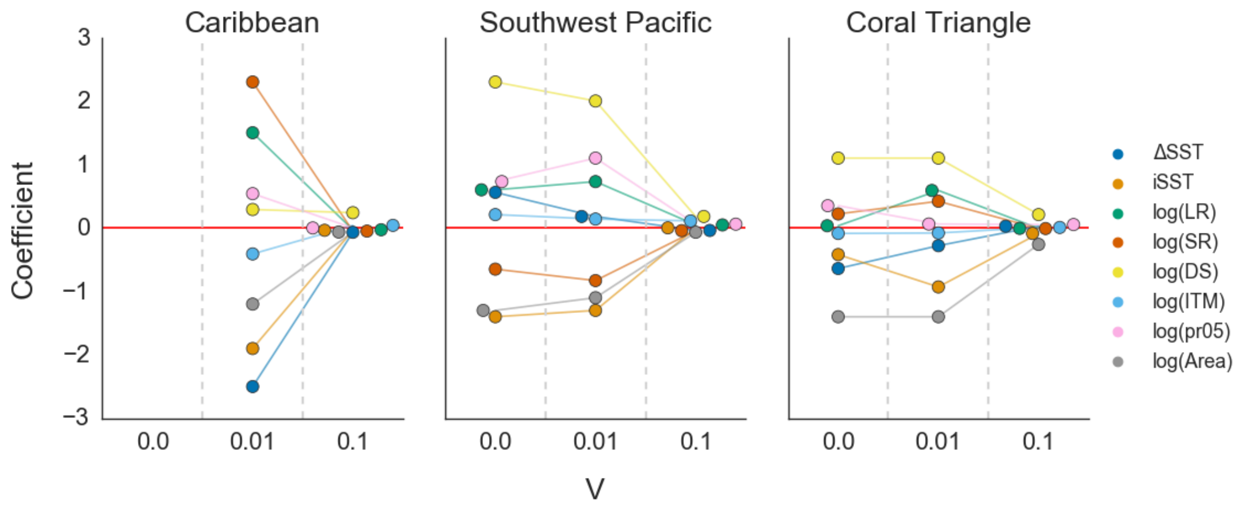


**Figure S9.** Standardized coefficient (effect size) of each covariate on RCP 8.5 minimum coral cover for *β*=0.5 with three levels of *V* in the Caribbean, Southwest Pacific and Coral Triangle. **Δ**SST is the change in sea surface temperature over the projection period, iSST is the initial temperature at the start of the projection, LR is local retention, SR is self-recruitment, DS is destination strength, ITM is initial temperature mismatch and pr05 is the proportion of incoming links from sites that are at least 0.5 °C warmer (based on iSST). Note that coefficients for the Caribbean model at *V*=0 are not shown because the GLM did not converge.

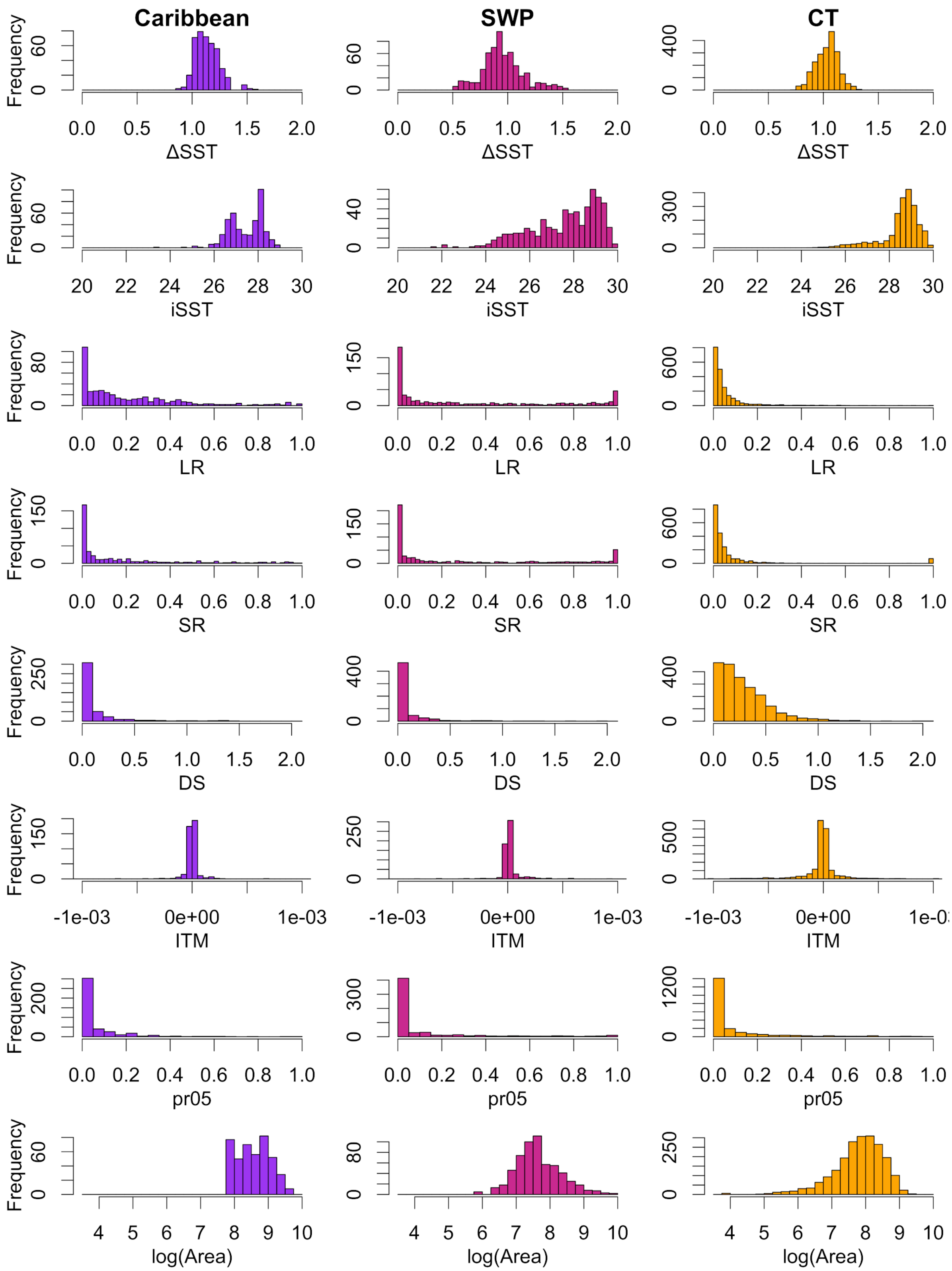


**Figure S10.** Distributions of covariates used in the GLMs for each region (purple for Caribbean, pink for SWP, and orange for CT).
